# Cellular and behavioral characterization of *Pcdh19* mutant mice: subtle molecular changes, increased exploratory behavior and an impact of social environment

**DOI:** 10.1101/2020.08.27.271239

**Authors:** Natalia Galindo-Riera, Sylvia Adriana Newbold, Monika Sledziowska, Jessica Griffiths, Erik Mire, Isabel Martinez-Garay

## Abstract

Mutations in the X-linked cell adhesion protein PCDH19 lead to seizures, cognitive impairment and other behavioral comorbidities when present in a mosaic pattern. Neither the molecular mechanisms underpinning this disorder, nor the function of PCDH19 itself are well understood. By combining RNA *in situ* hybridization with immunohistochemistry and analyzing single cell RNAseq datasets, we provide a first account of the subtypes of neurons expressing *Pcdh19*/*PCDH19*, both in the mouse and the human cortex. Our quantitative analysis of the *Pcdh19* mutant mouse reveals subtle changes in cortical layer composition, with no major alterations of the main axonal tracts. However, *Pcdh19* mutant animals, particularly females, display preweaning behavioral changes, including reduced anxiety and increased exploratory behavior. Our experiments also reveal an effect of the social environment on the behavior of wild-type littermates of *Pcdh19* mutant mice when compared with wild-type animals not housed with mutants. This is a second case of a mutated X-linked gene encoding a membrane protein expressed in the developing cortex impacting the behavior of co-housed wild-type littermates.

## INTRODUCTION

*PCDH19* is one of several genes located on the X chromosome known to impact neurodevelopment and behavior. Mutations in this gene were identified in patients suffering from EIEE9 (Epileptic Encephalopathy, Early Infantile, 9, OMIM #300088), also known as Girls Clustering Epilepsy (GCE), over a decade ago (Dibbens et al. 2008). Since then, more than 140 mutations have been described (Kolc et al. 2018), consolidating *PCDH19* as the second most relevant gene in epilepsy after *SCNA1* (Depienne and Leguern 2012; Duszyc et al. 2015). The pathogenicity of *PCDH19* mutations is dependent on cellular mosaicism and therefore the disorder follows an unusual inheritance, manifesting in heterozygous females and in males with somatic mutations (Depienne et al. 2009; Terracciano et al. 2016). Affected patients develop symptoms during early infancy, often within their first year of life, and display clustered seizures, varying degrees of cognitive impairment and other comorbidities, including autism spectrum disorder, attention deficits and obsessive-compulsive features (Kolc et al. 2020).

*PCDH19* codes for protocadherin 19, a calcium-dependent cell-cell adhesion molecule of the cadherin superfamily. This delta 2 protocadherin has 6 extracellular cadherin repeats, a single transmembrane domain and a cytoplasmic tail with two conserved motives of unknown function (CM1 and CM2, Wolverton and Lalande 2001). In addition, a WRC interacting receptor sequence (WIRS) downstream of CM2 allows PCDH19 to interact with the WAVE regulatory complex, enhancing its Rac1-mediated activation (Chen et al. 2014). PCDH19 is involved in different processes, ranging from neurulation and organization of the optic tectum in zebrafish (Emond et al. 2009; Cooper et al. 2015) to neurogenesis and regulation of GABAergic transmission in mammals (Fujitani et al. 2017; Bassani et al. 2018; Homan et al. 2018; Lv et al. 2019). In addition, PCDH19 is involved in gene expression regulation with estrogen receptor alpha (Pham et al. 2017) and mutations in *PCDH19* lead to a deficiency of the neurosteroid allopregnanolone and of other neuroactive steroids (Tan et al. 2015; Trivisano et al. 2017).

To date, two different *Pcdh19* knockout (KO) mouse models have been developed to explore the function of PCDH19. The first, produced by Taconic Biosciences, has the first three exons of the gene replaced by a beta galactosidase and neomycin (*LacZ-neo*) resistance cassette (Pederick et al. 2016). The second model retains exons 2 and 3, with a *Lacz-neo* selection cassette replacing exon 1, which encodes the entire extracellular and transmembrane domains (Hayashi et al. 2017). Lack of *Pcdh19* mRNA and protein was confirmed for the Taconic mouse (Pederick et al. 2016) and no major anatomical defects were reported in either of the two models. However, increased neuronal migration has been described (Pederick et al. 2016), as well as behavioral alterations (Hayashi et al. 2017; Lim et al. 2019). In addition, heterozygous females display a striking segregation of *Pcdh19* expressing and non-expressing progenitors in the developing cortex and altered electrocorticogram traces (Pederick et al. 2018).

Although no major abnormalities in cortical architecture have been reported in either KO mouse model, no detailed, quantitative analysis has been carried out yet. Similarly, while RNA *in situ* hybridization revealed strongest *Pcdh19* expression in layers II/III and V(a) in mice (Pederick et al. 2016; Hayashi et al. 2017), the neuronal subtypes expressing *Pcdh19* have not been characterized, possibly due to the difficulty of labeling PCDH19 expressing cells with current antibodies. Here we report on the identity of *Pcdh19* expressing neurons in the mouse and human cortex. We also uncover subtle alterations in cortical neuronal distribution in the cortex of Taconic *Pcdh19* mutant animals, as well as robust differences in the behavior of heterozygous females, including an impact of mutant animals on the behavior of their wild type littermates.

## MATERIAL AND METHODS

### Experimental Animals

Animals were housed under a 12h light/dark cycle with *ad libitum* access to water and food, and controlled temperature and humidity. All experiments using mice were conducted at Cardiff University, approved by the local ethical boards and carried out following the directions of the UK Animal Scientific Procedures Act (update 1986).

C57BL6/J wild-type (WT) animals were purchased from Charles River Laboratories and the *Pcdh19* knock-out (KO) line (TF2108) was acquired from Taconic Biosciences. Experimental matings for anatomical and cellular characterization, as well as for behavioral analysis were set up using wild type males and *Pcdh19* heterozygous (HET) females to produce litters with WT males and females, KO males and HET females.

### Analysis of single cell RNAseq datasets

Gene expression matrices and metadata were downloaded from https://portal.brain-map.org/atlases-and-data/rnaseq. Analysis and visualization were carried out using R v.3.6.3, assisted by RStudio v.1.2.1335. Raw counts were normalised to account for library size (total sum of counts per cell) and transformed to counts per million (CPM) using R package scater v.1.16.2. Violin plots were generated with R packages gridExtra v.2.3 and ggplot2 v.3.3.1. River plots were made with R packages gridExtra v.2.3, ggplot2 v.3.3.1 and ggforce v.0.3.2.

### Tissue Processing

Animals were perfused with PBS followed by 4% paraformaldehyde (PFA) in PBS. After perfusion, brains were extracted and post-fixed in PFA 4% overnight at 4 C. For RNA ISH, brains were then cryoprotected in 30% sucrose in PBS before embedding in OCT compound (Tissue-Tek) prior to freezing. Samples were stored at −80 C until sectioning. 12 or 20 μm sections were cut with a cryostat (CM3050, Leica Systems) and stored at −80 C until use. For immunostaining, fixed brains were briefly washed in PBS and embedded in 4% low melting point agarose. 50 μm sections were cut with a vibrating microtome (VT1000S, Leica Systems) and stored in PBS with 0.05% Sodium Azide at 4 C until use.

### RNA *in situ* Hybridization and Immunohistochemistry

The probe to detect *Pcdh19* has been described before (Gaitan and Bouchard 2006). Its sequence was amplified using primers Pcdh19e1-F, 5’-CACCAAGCAGAAGATTGACCGAG-3’ and Pcdh19e1-R, 5’-GCCTCCCATCCACAAGAATAGTG-3’ and cloned into pCRII-Blunt-TOPO (Invitrogen). This plasmid was then used to generate digoxigenin-labeled sense and antisense probes.

Thawed sections were post-fixed in 4% PFA, endogenous peroxidases were quenched with 3% hydrogen peroxidase and slices were then acetylated in a 0.25% acetic anhydride solution. Pre-hybridization took place in pre-warmed hybridization buffer (50% formamide, 0.1% Tween-20, 0.25% CHAPS, 250 μg/ml yeast tRNA, 500 μg/ml herring sperm, 5x Denhardts, 5x SSC, 50 μg/ml heparin, 2.5 mM EDTA) for 1h at 65 C. Slices were hybridized with the denatured sense or antisense probes overnight at 65 C in a humidified chamber. The next day, slides were washed with 0.2x SSC (GIBCO) and PBST, and then blocked in ISH blocking solution (10% DS and 0.1% TritonX-100 in PBS) for 20 min at RT. After blocking, brain slices were incubated in primary antibody for 1 h at RT; washed in PBST and incubated in secondary antibody for 1 h at RT. Antibodies used are described below. Slides were then washed in PBST, equilibrated in TN buffer (150 mM NaCl and 100 mM Tris pH= 7.5 in water) and incubated for 30 min in 1:2000 HRP-coupled anti-DIG antibody (Sigma-Aldrich, 11207733910). Following the incubation, tissue was rinsed in TNT (TN + 0.5% Tween) and immersed in Cy3-Tyramide (TSATM Plus Cy3 Fluorescence kit, Perkin-Elmer, NEL744001KT) in a 1 in 50 dilution dissolved in the amplification diluent. Slides were then washed, counterstained with DAPI and mounted in DAKO.

### Immunohistochemistry

Antigen retrieval was performed for stainings with antibodies against RORB, SATB2, Pvalb and CR, with the tissue either immersed in a 10 mM citrate buffer pH =6, at 95 C for 5 min (RORB and SATB2) or 10 min (Pvalb, CR) before blocking. 50 μm coronal sections were blocked (4% BSA, 3% donkey serum, 0.1% Triton X-100 in PBS) at RT for 1 h. The tissue was then incubated in primary antibody diluted in blocking solution overnight at 4 C. Primary antibodies used for immunostaining were as follows: anti-CUX1 rabbit polyclonal (1:200; Proteintech, 11733 or Santa Cruz Biotechnology, sc-13024), anti-CTIP2 rat monoclonal (1:250; Abcam, ab18465), anti-SATB2 mouse monoclonal (1:400; Abcam, ab51502), anti-RORB rabbit polyclonal (1:200; Proteintech, 17635-1AP), anti-TBR1 rabbit polyclonal (1:350; Abcam, ab31940), anti-Pvalb rabbit polyclonal (1:10000 or 1:500 for ISH; Swant, PV27), anti-CB rabbit polyclonal (1:5000; Swant, CB38), anti-CR mouse polyclonal (1:1000; Merck, AB5054), anti-SST rat monoclonal (1:200; Merck, MAB354), anti-L1CAM rat monoclonal (1:500, Merck, MAB5272), anti-Neuropilin1 goat polyclonal (1:300, R&D Systems, AF566).

Slices were then rinsed in PBS and incubated with secondary antibodies coupled to fluorochromes (Alexa Fluor range, Thermo Fisher Scientific) for 1 h at RT. Nuclei were counterstained with DAPI for 10 min, washed again in PBS and mounted with DAKO mounting medium.

### Image acquisition and analysis

Images were acquired using a confocal laser scanning microscope (LSM 780, Carl Zeiss) and ZEN Black software (version 2.0, Carl Zeiss). Image analysis was conducted with ImageJ Fiji software (Schindelin et al. 2012). For quantification, the cortical wall was divided into ten horizontal bins of equal width. The number of marker positive cells in each bin was quantified and is shown as mean percentage relative to the total number of cells in all ten bins, ± standard error of the mean (SEM).

### Behavioral analysis

Behavioral tests were conducted at P21 (pre-weaning) and in young adults (P60 and over). Two different WT controls were tested: WT littermates of the mutant animals (mixed genotyped housing mice, MGH) and animals from pure WT litters (single genotype housed mice, SGH). The WT parents of the SGH animals were derived from the *Pcdh19* colony. Mice habituated to the new environment by taking them to the behavioral room 30 min prior to the tests. Mice were handled with open hands to reduce anxiety levels and a maximum of one behavioral test was performed per day.

### Open field

Open field behavioral analysis was performed on two consecutive days, using the first day to habituate the mice to the new environment. Mice were allowed to explore freely, in the dark, for 20 min, in an open field arena (40 cm x 40 cm). Spontaneous locomotion was recorded using a computer-linked video camera (The Imaging Source) located above the arena and an infrared illumination box (Tracksys) located underneath the arena. The EthoVision XT software (Noldus) was used to analyze total distance travelled, distance travelled in intervals of 5 min and time spent in the center of the arena. The center of the arena was defined as the area separated from the wall by 5 cm or more.

### Elevated plus maze

Each mouse was left to explore freely for 5 min in a maze consisting of 4 perpendicular arms (40 cm x 7 cm): two open arms (1 cm high) and two closed arms (16 cm high), in a well-lit room. Behavior was recorded using a computer-linked video camera (The Imaging Source) located above the maze. Total time spent in the open arms was measured using EthoVision XT software.

### Social interaction

At P21, test pups were habituated to the arena for 3 min. Subsequently, WT females in estrous, unfamiliar to the pups, were added to the cage and both mice were allowed to interact with each other for another 3 min in a well-lit room. The interaction between the pups and the females was recorded using a computer-linked video camera (The Imaging Source) located above the arena. Videos were manually scored, and interaction recorded when both mice were within 2 cm of each other, not including tail-tail interactions. At P60, only female mice were tested for social interaction. In this case the unknown WT females were not required to be in oestrus.

To determine which females were in oestrus, vaginal smears were stained with Giemsa solution (Polysciences inc.) (Caligioni 2009) prior to the experiment.

### 24-hour activity

P60 experimental mice were placed in individual clear boxes (40 cm × 24 cm × 18 cm) and let to roam free for 24 h with *ad libitum* access to food and water and their normal 12 h light/dark cycle. Three infrared beams traversed each cage at the bottom. Data were analyzed using the MED-PC® IV software suite and extracted using the MPC2XL program. The number of beams breaks in 24 h and in 1-hour slots, as well as the total number of beam breaks during the light and dark periods were analyzed.

### Experimental design and Statistical analysis

For all experiments, individual animals were considered the experimental unit and data obtained from each animal was averaged if more than one quantification was performed (for example when analysing several brain slices from the same animal). Experimenters were blind to the genotype of the animals until all quantification or scoring was completed. Statistical analysis was performed using IBM SPSS Statistics® 25 software (cortical lamination analysis) or R software (behavior), version 3.6.2. (R Core Team 2019). Normality of the data was tested using the Shapiro-Wilk test and homogeneity of variance was assessed with Levene’s test. If either assumption was violated an appropriate non-parametric test was used. For comparison of more than two groups, analysis of variance (ANOVA) was used for normal data and Kruskal-Wallis if the assumption of normality was not met. If only the assumption of homogeneity of variance was not met, a Welch’s ANOVA test was used. Post-hoc test following ANOVA was adjusted according to Tukey’s HSD or, in the case of the social interaction analysis, Dunnet’s test. Kruskal-Wallis was followed by Dunn’s correction and Welch’s ANOVA was followed by Games-Howell correction. All statistical data are presented as mean ± SEM.

## RESULTS

### *Pcdh19* is expressed by different subtypes of cortical projection neurons and interneurons

Previous RNA *in situ* hybridization (ISH) studies have shown two main areas of *Pcdh19* expression in the cortex, corresponding to the upper regions of layer V (layer Va) and II/III (Hertel and Redies 2010; Pederick et al. 2016). However, a detailed analysis of the cortical neuronal subtypes expressing *Pcdh19*, an important consideration given the cellular diversity of the cortex, is still lacking. ISH against *Pcdh19* was combined with immunohistochemistry (IHC) against several cortical markers for principal neurons and interneurons in the somatosensory cortex at postnatal days 10 (P10) and P20, respectively (Fig. 1 A-D). At P10, *Pcdh19*+ cells were found to co-express markers for layer IV neurons (RORB, Fig. 1A), callosal projection neurons (SATB2, Fig. 1B), corticospinal neurons (CTIP2, Fig. 1B), and corticothalamic neurons (TBR1, Fig. 1C). Strongest co-expression was seen in SATB2+ neurons, whereas RORB+ cells showed weaker expression and in a smaller proportion of cells. CTIP2+ neurons with strong *Pcdh19* expression tended to be located in the upper half of layer V, whereas TBR1+ cells co-expressing *Pcdh19* did so at generally lower levels. At P20, interneurons were identified co-expressing *Pcdh19* with Parvalbumin in layers II/III and V (Fig. 1D), as well as double positive cells for Calbindin and *Pcdh19* (data not shown). This approach is limited to available antibodies that withstand the harsh treatments needed for ISH. In addition, recent studies using single cell transcriptomics on mouse and human cortical tissue have revealed numerous subtypes of excitatory and inhibitory neurons that cannot be readily identified through immunostaining. In view of these limitations, we then turned to publicly available datasets of cortical single cell RNA expression and re-analyzed them to assess *Pcdh19* expression in the mouse adult cortex (Fig. 1 E, F). We chose the dataset published by Tasic *et al* in 2018 (Tasic et al. 2018), which includes 23,822 single-cell transcriptomes with cluster-assigned identity isolated from the primary visual (VISp) and anterior lateral motor (ALM) cortices of adult mice (Mouse – V1 & ALM – SMART-seq, available at https://portal.brain-map.org/atlases-and-data/rnaseq). This study currently provides the most comprehensive classification of cortical cellular types to date with 133 cell types, of which 56 are glutamatergic, 61 GABAergic and 16 non-neuronal. In agreement with our P10 and P20 results, scRNAseq data shows that *Pcdh19* expression is maintained in both excitatory and interneuronal cortical populations in the adult that co-express the markers of our ISH analysis (Fig. 1E, F).

**Figure 1.**
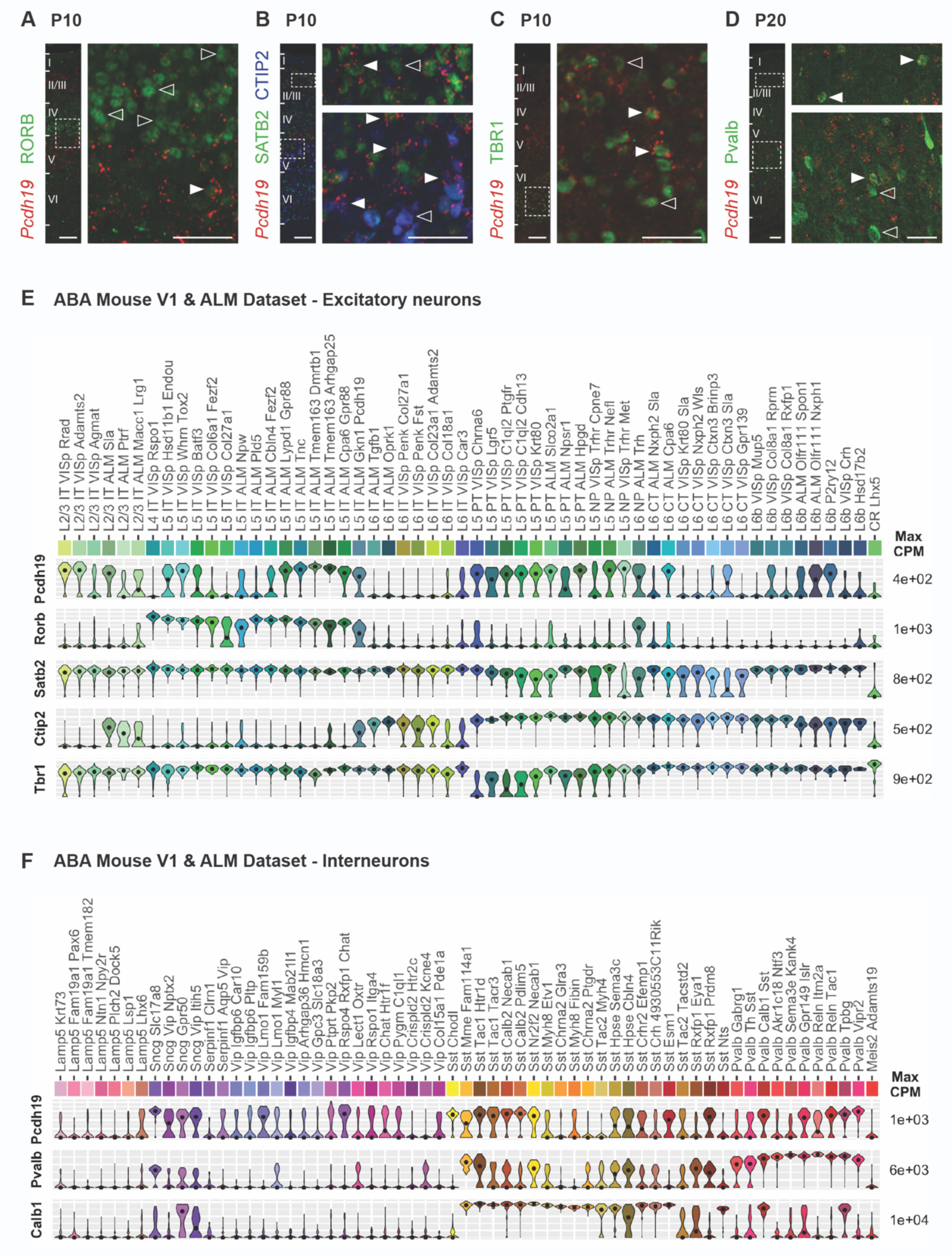
*Pcdh19* is expressed by excitatory and inhibitory neurons in the mouse cortex. (A-D) Confocal micrographs of P10 (A-C) and P20 (D) cortical slices hybridized with an RNA probe against *Pcdh19* (red) and antibodies against (A) RORB (green), (B) SATB2 and CTIP2 (green and blue, respectively), (C) TBR1 (green) and (D) Parvalbumin (Pvalb, green). The left panel shows the entire cortical wall with boxes indicating the regions enlarged in the right panels. White arrowheads point to double positive cells, empty arrowheads point to single positive cells (*Pcdh19* negative). Scale bars: left panels, 100 μm; right panels, 50 μm. (E, F) Violin plots representing gene expression and distribution for *Pcdh19* and the markers used in (A-D) in the different excitatory (E) and interneuronal (F) subtypes defined in the Tasic et al 2018 study (Allen Brain Atlas Mouse V1 & ALM Dataset). Dots indicate the median value of the cluster in CPM. CPM values are displayed on log10 scale.

*Pcdh19* expression is cluster-specific in all layers and, in general terms, within projection neurons *Pcdh19* expression is most consistent in layer V cells, particularly in those projecting through the pyramidal tract (PT) and in the newly defined near-projecting population (NP) (Fig. 1E). Intratelencephalically projecting neurons in layer V show lower expression levels in the primary visual cortex, but much higher levels in the anterior motor cortex, with the two L5 IT ALM Tmem163 subpopulations (Dmrtb1 and Arhgap25) displaying the strongest expression of all characterized neuronal subtypes. Interestingly, one particular L5 IT ALM cluster is defined by *Pcdh19* expression (L5 IT ALM Gkn1 Pcdh19), although it does not exhibit the highest expression levels for this gene and is also not included in any of the homology clusters with human neurons (see below). Expression in layer II/III is lower on average, but some neuronal clusters (L2/3 IT VISp Rrad, L2/3 IT VISp Adamts2 and L2/3 ALM Sla) present equivalent expression levels as layer V cells, both in the visual and the motor cortex. Expression of *Pcdh19* in layer VI neurons is negligible in the visual cortex, but stronger in corticothalamic and near-projecting cells in the anterior motor cortex. Expression in layer VIb follows a similar pattern (Fig. 1E). Therefore, expression of *Pcdh19* in excitatory neurons shows regional differences between brain areas, with generally higher expression in the anterior lateral motor cortex than in the primary visual cortex.

As in the case of projection neurons, *Pcdh19* expression in interneurons is strongly cluster-dependent. More specifically, strongest average expression is found in the *Sncg* and *Pvalb* subclasses and the *Sst-Chodl* type; however, there is considerable variation and several *Sst* clusters also express *Pcdh19* widely (Fig. 1F). *Sncg* neurons are *Vip*+, *Cck*+ multipolar or basket cells located mainly in upper layers, and 3 out of their 4 subtypes have consistent *Pcdh19* expression. Only one other cluster of *Vip* interneurons, corresponding to bipolar or multipolar cells, also shows relevant *Pcdh19* expression (Vip Rspo4 Rxfp1 Chat). Within the *Pvalb* subclass, *Pcdh19* is expressed by Chandelier cells (Pvalb Vipr2) and several subtypes of basket cells (Pvalb Gpr149 Islr, Pvalb Reln Tac1 and Pvalb Tpbg). Finally, within the *Sst* subclass, *Pcdh19* expression is strongest in some subtypes of upper layer basket cells (Sst Tac1 Htr1d, Sst Tac1 Tacr3 and Sst Nr2f2 Necab1) and Martinotti cells (Sst Calb2 Necab1 and Sst Calb2 Pdlim5), and in the long-range projecting population (*Sst-Chodl*).

In summary, our analysis demonstrates that mouse *Pcdh19* is expressed by a heterogeneous neuronal population that includes a wide variety of cortical projection neurons and interneurons, with strong variation between neuronal subtypes. Expression in non-neuronal cells is very low (data not shown).

### Human *PCDH19* is also expressed in excitatory and inhibitory neurons

The expression profile of human *PCDH19* and a comparison with its murine counterpart is relevant with regards to the use of *Pcdh19* mutant mice as a model to investigate the pathophysiology of the disorder caused by mutations in the human gene. Human cortical neurons have also been characterized at the transcriptional level using single cell RNAseq (Lake et al. 2016; Hodge et al. 2019). We therefore chose to analyze a dataset obtained from predominantly neuronal nuclei of the human middle temporal gyrus (MTG; Human – MTG – SMART-seq, available at https://portal.brain-map.org/atlases-and-data/rnaseq) because it has been systematically compared to the mouse data described above (Hodge et al 2019). That comparison yielded 32 homologous neuronal clusters, allowing a meaningful comparison of the expression of *PCDH19* between species (Suppl. Fig.1). In this human dataset, *PCDH19* expression in excitatory neurons is more prevalent in layer V neurons projecting through the pyramidal tract, whilst all other excitatory clusters show lower expression levels (Fig. 2A). In contrast, different interneuronal types display strong *PCDH19* expression, including several VIP and Parvalbumin clusters, as well as Chandelier cells (Fig. 2A). When expression is compared between species across the 32 homology clusters, only Chandelier cells and layer V PT projecting neurons show strong expression in both human and mouse (Fig. 2B). In addition, there is also strong expression in subpopulations of bipolar or multipolar interneurons in both species (cluster VIP4, mouse subtype Vip Rspo4 Rxtp1 Chat; human subtype Inh L1-3 VIP-GGH), although the pooling of several subtypes into cluster VIP4 dilutes the overall expression. In general, *PCDH19* expression in human neurons seems to be restricted to fewer cell types, while in the mouse, higher levels of expression are found in most excitatory clusters and in inhibitory clusters Pvalb2, Sst2, Sst3, Sst4, Sst5, Sst Chodl and Vip Sncg. In human, neurons in clusters Lamp5 Lhx6 and VIP2 show higher expression levels than their murine counterparts.

**Figure 2.**
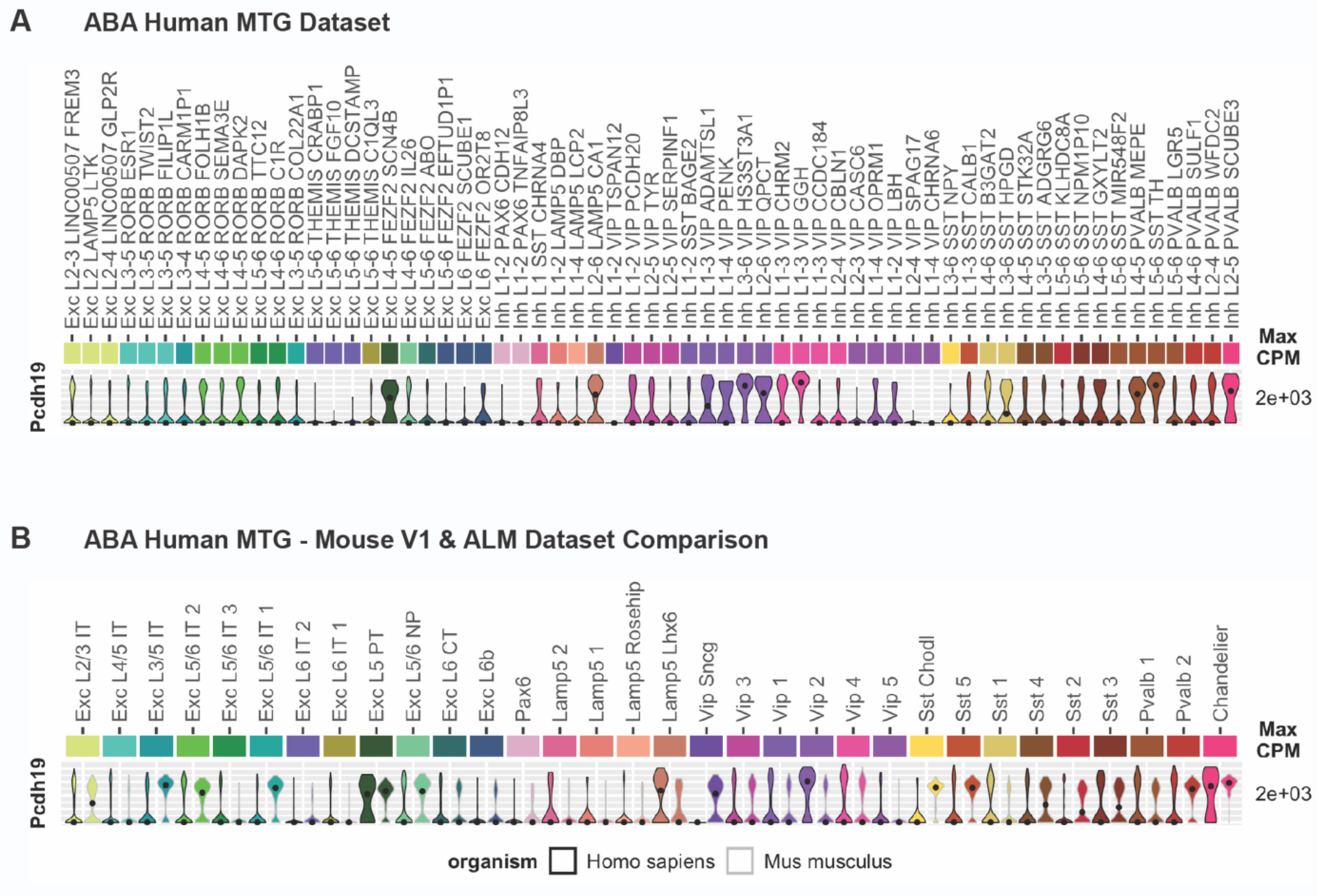
*PCDH19* is expressed by excitatory and inhibitory neurons in the human cortex. (A) Gene expression and distribution of *PCDH19* in the Allen Brain Atlas human MTG dataset, represented by violin plots. (B) Comparison of gene expression and distribution between mouse and human neurons in the 32 homology clusters defined by Hodge et al in their 2019 study. For each cluster, human data are shown on the left (black outline) and mouse data on the right (grey outline). Dots indicate the median value of the cluster in CPM. CPM values are displayed on log10 scale.

As the mouse data presented above show that *Pcdh19* expression vary between brain regions (anterior lateral cortex and primary visual cortex in this case), we extended our analysis to a second human dataset from the Allen Brain Atlas, obtained from several different brain areas (middle temporal gyrus, anterior cingulate gyrus, primary visual cortex, primary motor cortex, primary somatosensory cortex and primary auditory cortex) that is also publicly available (Human – Multiple Cortical Areas – SMART-seq, available at https://portal.brain-map.org/atlases-and-data/rnaseq). This dataset, which also includes the MTG data, comprises 49,417 cell nuclei, compared with 15,928 for the middle temporal gyrus alone. It has therefore allowed the definition of 56 excitatory and 54 inhibitory subtypes, as opposed to 23 and 41, respectively, in the MTG dataset (Suppl. Fig. 2 shows the correspondence between the clusters in the two datasets). In addition to analyzing expression levels across the whole dataset, using the metadata of individual nuclei, we separated the expression of *PCDH19* in the different excitatory and inhibitory clusters by brain region (Suppl. Fig. 3 and 4). Our analysis confirmed that, as a whole, within excitatory neurons strongest and most consistent *PCDH19* expression is found in FEZF2 subtypes in layer V or layers III-V, but it also revealed strong expression of *PCDH19* in several other excitatory neuronal subtypes spanning layers II-V, particularly Exc L3 RORB CARTPT, Exc L3-4 RORB FOLH1B, Exc L4-5 RORB LCN15 and Exc L5 RORB SNHG7 (Suppl. Fig. 3). In addition, low expression is evident in many other excitatory neurons of layers III-VI, although several layer IV clusters, particularly in sensory regions, tend to express much lower levels of PCDH19, as was the case in the mouse. Regarding interneurons, *PCDH19* expression is highest in the L3-6 VIP KCTD13 subtype, with strong expression in all cells. In addition, *PCDH19* is also relatively highly expressed in several other VIP, LAMP5, SST and PVALB subpopulations (Suppl. Fig. 4). A comparison between different brain regions reveals that, in general, *PCDH19* is expressed in each cluster at similar levels across areas. However, there are some exceptions, like L1 VIP PCDH20 interneurons, which show much higher *PCDH19* expression in the visual cortex (V1C) than in somatosensory areas (S1lm and S1ul) or L1-2 VIP RPL41P3, with higher *PCDH19* expression in motor areas (Suppl. Fig. 4).

**Figure 3.**
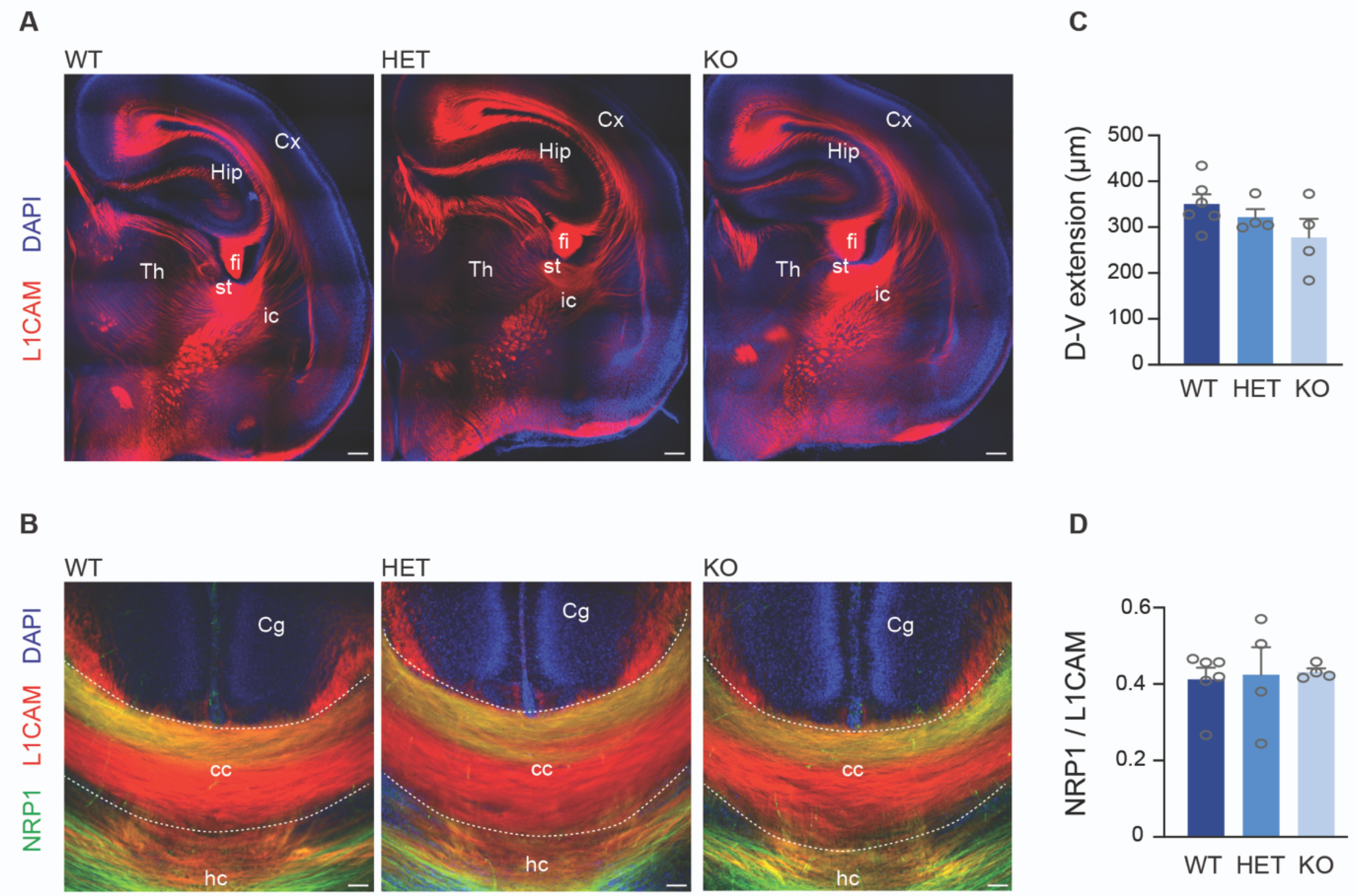
No major anomalies in the main axonal tracts in *Pcdh19* mouse mutants. (A) Confocal micrographs of P0-P1 mouse hemispheres stained with anti-L1CAM (red). Nuclei were counterstained with DAPI (blue). (B) Confocal micrographs of the corpus callosum of P0-P1 mice stained with anti-L1CAM (red), anti-Neuropilin-1 (green) and counterstained with DAPI (blue). (C) Quantification of the dorso-ventral extension of the corpus callosum in WT and mutant animals. (D) Quantification of the dorsal restriction of Neuropilin-1 positive axons in WT and mutant animals. All results are indicated as mean ± SEM. 2 images per brain, obtained from four animals originating from three different litters were analyzed for each condition. Cx, cortex; Hip, hippocampus; Th, thalamus, fi, fimbria; st, striatum; ic, internal capsule; Cg, cingulate cortex; cc, corpus callosum; hc, hippocampal commissure. Scale bars: 200 µm (A) and 50 µm (B).

**Figure 4.**
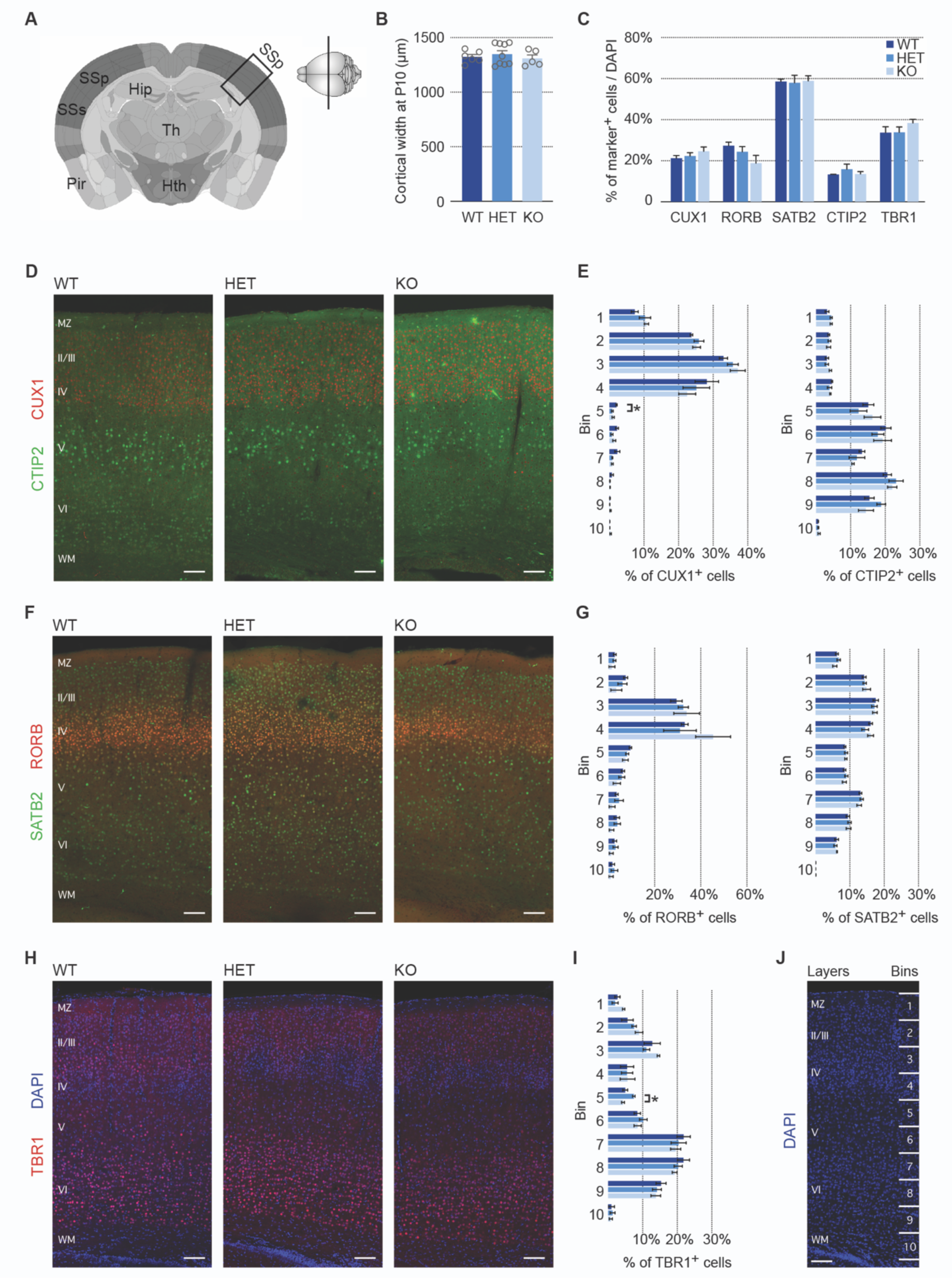
Minor changes in the distribution of cortical excitatory neurons in *Pcdh19* mutant animals. (A) Diagram indicating the position of the cortical area analyzed (primary somatosensory cortex, SSp). (B) Quantification of cortical width at P10 in *Pcdh19* WT, HET and KO animals. (C) Relative percentages of the different cortical markers examined with respect to total DAPI+ cells. (D) Representative confocal micrographs of immunohistochemistry with anti-CUX1 (red) and anti-CTIP2 (green) antibodies on WT, HET and KO tissue. (E) Quantification of the percentage of marker+ cells in each of 10 equal bins spanning the cortical wall. (F) Representative confocal micrographs of immunohistochemistry with anti-RORB (red) and anti-SATB2 (green) antibodies on WT, HET and KO tissue. (G) Distribution of marker+ cells in each of 10 equal bins spanning the cortical wall, shown as percentage. (H) Representative confocal micrographs of immunohistochemistry with anti-TBR1 (red) antibodies on WT, HET and KO tissue. Nuclei are counterstained with DAPI (blue). (I) Quantification of the percentage of marker+ cells in each of 10 equal bins spanning the cortical wall. (J) Confocal micrograph of the SSp counterstained with DAPI (blue), indicating the correspondence between cortical layers and the bins used for quantification. All results are indicated as mean ± SEM. A minimum of 3 images per brain, obtained from four animals originating from three different litters were analyzed for each condition. *P < 0.05. Scale bars: 200 μm.

Overall, although in a direct comparison of the 32 homology clusters *PCDH19* expression seemed to be more widespread in mouse than in human neurons, this was probably due to the particular brain region sampled. The larger and more diverse human sample allows a more fine-grained neuronal classification, revealing more prevalent expression of *PCDH19* among excitatory and inhibitory neurons and suggesting some regional variations between cortical areas in the human brain, as seen also in the mouse brain.

### No obvious defects in axonal tracts in *Pcdh19* mutant animals

Our results indicate that *Pcdh19* is expressed in cortical projection neurons that project through the corpus callosum (layer II-III and some layer V neurons), as well as in neurons projecting outside the cortex, mainly through the pyramidal tract (layer V PT neurons). Although several members of the cadherin superfamily, including delta protocadherins 7, 10, 17 and 18, have been shown to play a role in axonal outgrowth (Uemura et al. 2007; Piper et al. 2008; Hayashi et al. 2014), fasciculation (Williams et al. 2011; Hayashi et al. 2014) and arborization (Biswas et al. 2014), it is not known whether mutations in *Pcdh19* have an impact on any of these processes. We therefore conducted a general characterization of axonal tracts in Taconic *Pcdh19* WT, KO and HET animals by immunostaining against the cell adhesion molecule L1CAM (Fig. 3A). No differences were apparent between genotypes in the major axonal tracts, including the internal capsule, stria terminalis, fimbria or corpus callosum. Next, we analyzed the corpus callosum in more detail by labelling dorsally located axons with Neuropilin-1, which allows the analysis of topographical organization at the midline. Again, the dorso-ventral extension of the corpus callosum and the dorsal restriction of Neuropilin-1 expressing axons was similar between all genotypes (Fig. 3B-D) (D-V extension: Kruskal-Wallis, *H* = 2.86, *P* = 0.2635, df = 2; dorsal restriction: Kruskal-Wallis, *H* = 0.07, *P* = 0.9783, df = 2). Thus, our results revealed no major abnormalities in the main axonal tracts, although they do not preclude the existence of more subtle defects that would require a more detailed analysis to be revealed.

### Cortical lamination is largely preserved in *Pcdh19* mutant animals

Although no major lamination defects have been described in *Pcdh19* mutant animals (Pederick et al. 2016; Hayashi et al. 2017), no detailed quantitative studies have been performed so far that could reveal subtle changes in layer composition. Given that *Pcdh19* is expressed in projection neurons and interneurons, we performed an analysis with markers for both neuronal populations. We first selected cortical markers for different types of projection neurons (CUX1, SATB2, RORB, CTIP2 and TBR1) and performed immunohistochemistry in the somatosensory cortex (SSC) at P10, once radial migration is completed (Fig. 4A). For each marker, we determined the proportion of positive cells, as well as their distribution within 10 bins covering the whole width of the cortical plate (Fig. 4J). In accordance with previous reports (Pederick et al. 2016), we found no differences in cortical width between genotypes (WT = 1321.09 ± 21.59 μm, HET = 1354.23 ± 33.78 μm and KO = 1305.41 ± 30.62 μm; Fig. 4B). The proportion of positive neurons for all five examined markers was also unaltered (Fig. 4C). CUX1+ cells made up approximately one fifth of all DAPI+ cells (WT = 21.45 ± 1.2%, HET = 22.36 ± 1.65%, KO = 24.82 ± 2.13%) and SATB2+ cells represented more than half of all cells (WT = 59.08 ± 0.92%, HET = 58.08 ± 3.62%, KO = 57.82 ± 2.74%). The proportion of RORB+ cells seemed lower in KO brains, especially when compared to WT (WT = 27.73 ± 1.75%, HET = 24.43 ±2.5%, KO = 18.98 ± 3.7%), but statistical analysis revealed this difference was not significant (one way ANOVA, main effect of genotype *F*(2,9) = 2.39, *P* = 0.1468). CTIP2+ cells represented about 13% in all three genotypes (WT = 13.33 ± 0.46%, HET = 13.82 ± 1.32%, KO = 13.61 ± 1.14%) and TBR1+ cells added up to approximately one third of all cells (WT = 33.65 ± 2.92%, HET = 34.03 ± 2.64%, KO = 38.51 ± 1.7%). Furthermore, the distribution of SATB2+, RORB+ and CTIP2+ neurons between the 10 bins was unchanged (Fig. 4D-G). However, we detected some subtle deviations in the distribution of CUX1+ and TBR1+ neurons, in both cases regarding bin 5 (Fig. 4D, E, H-J). PCDH19-HET animals showed a significant 2.4-fold reduction in the percentage of CUX1+ neurons in this bin compared to wild types (WT = 2.08 ± 0.18%, HET = 0.86 ± 0.27%, KO = 1.14 ± 0.32%; one way ANOVA, main effect of genotype *F*(2,9) = 5.81, *P* = 0.0239; Tukey: *q*(1,9) = 4.60, *P* = 0.0245 HET vs WT). Conversely, the percentage of TBR1+ cells in this bin was increased in HET brains when compared to KO brains (WT = 4.97 ± 0.71%, HET = 7.46 ± 0.35% and KO = 4.34 ± 0.41%; Kruskal-Wallis, *H* = 8.0, *P* = 0.0048, df = 2; post-hoc Dunn: *Z* = 2.75, *P* = 0.0181 HET vs KO).

To complete our analysis on cortical composition and lamination, we stained the SSC with four different interneuronal markers (SST, PVALB, CB, and CR) in P20 brains. As before, cortical thickness showed no difference between genotypes (WT = 1457.18 ± 32.71 μm, HET = 1402.97 ± 42.92 μm and KO = 1387.02 ± 9.88 μm; Fig. 5A) and we did not detect any significant changes in the proportion of marker positive neurons (Fig. 5B). The most abundant type was CB+ cells (WT = 19.11 ± 1.04%, HET = 16.20 ± 1.21%, KO = 18.77 ± 0.20%), whereas PVALB+, SST+ and CR+ accounted for less than 5% of DAPI+ cells each (PVALB: WT = 2.99 ± 0.28%, HET = 3.7 ± 0.2%, KO = 3.15 ± 0.21%; SST: WT = 2.22 ± 0.24%, HET = 1.33 ± 0.11%, KO = 1.61 ± 0.33%; CR: WT = 1.61 ± 0.42%, HET = 1.14 ± 0.04%, KO = 0.98 ± 0.1%). Because the absolute number of neurons counted for those three markers in the SSC was very low, we extended our analysis to include the adjacent motor cortex and compared the number of neurons positive for each marker, rather than the percentage over DAPI. No differences in the number of neurons were found in this analysis (Fig. 5C). Regarding cellular distribution in the SSC, no differences were apparent for CR+ and PVALB+ cells. However, we detected small but significant changes in the distribution of CB+ and SST+ interneurons between HET and KO brains (Fig. 5D-G). HET brains showed a 1.5-fold increase in the percentage of CB+ cells in bin 7 when compared to KO brains (WT = 2.17 ± 0.17%, HET = 2.95 ± 0.26%, KO = 1.98 ± 0.26%; one way ANOVA, main effect of genotype *F*(2,9) = 5.04, *P* = 0.034; Tukey: *q*(1,9) = 4.23, *P* = 0.0366 HET vs KO), and KO brains displayed a 1.7-fold increase over HET brains in the percentage of SST+ cells in bin 5 (WT = 11.21 ± 1.34%, HET = 10.58 ± 1.62%, KO = 17.71 ± 2.12; one way ANOVA, main effect of genotype *F*(2,9) = 5.26, *P* = 0.0307; Tukey: *q*(1,9) = 4.14, *P* = 0.0403 HET vs KO). In summary, the relative neuronal proportions and distribution are mostly normal in the SSC of *Pcdh19* mutant animals, although subtle differences in distribution cannot be ruled out for particular neuronal types.

**Figure 5.**
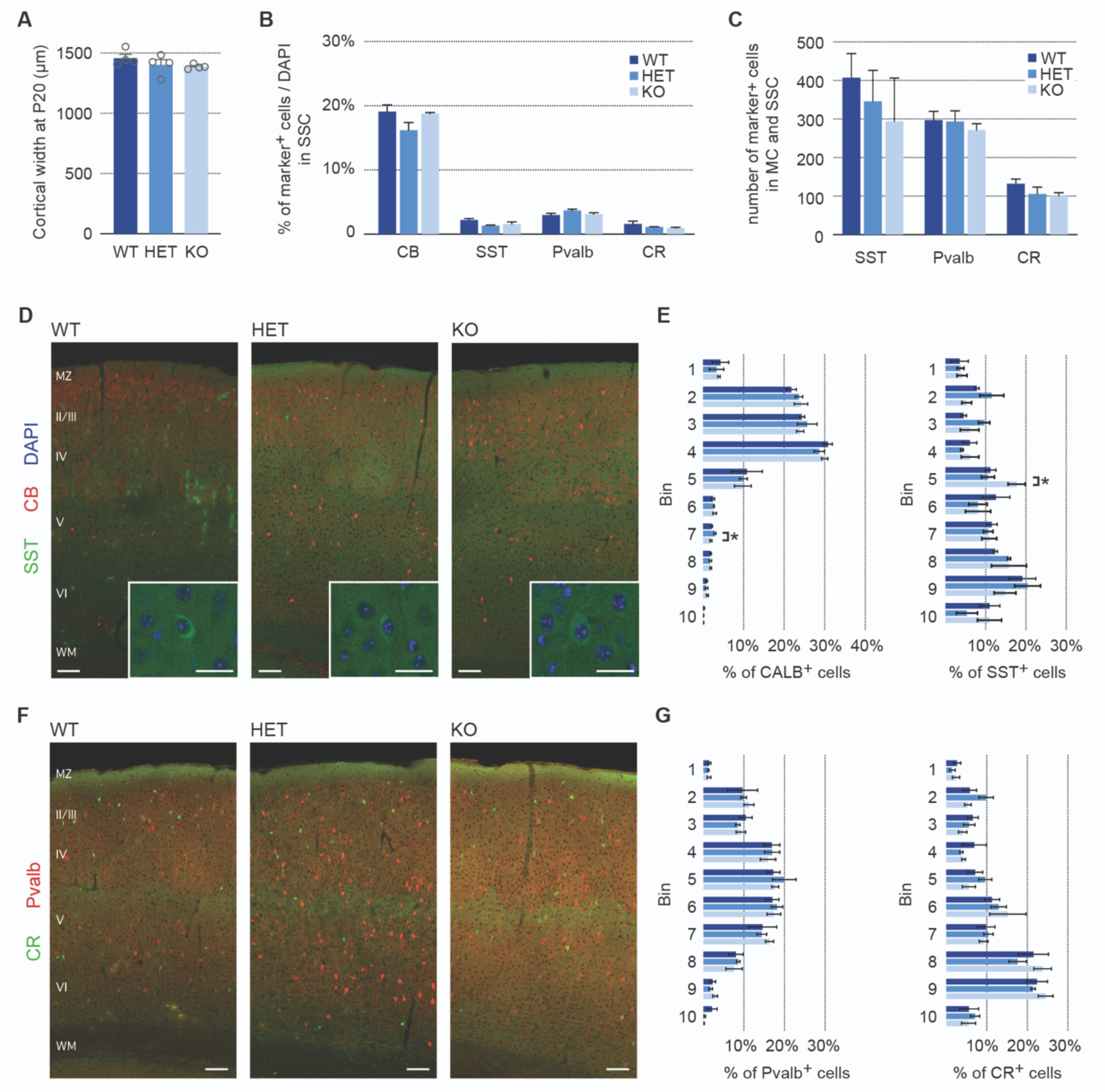
Subtle changes in the distribution of inhibitory neurons in the cortex of *Pcdh19* mutant animals. (A) Quantification of cortical width at P20 in *Pcdh19* WT, HET and KO animals. (B) Relative percentages of the different cortical markers examined with respect to total DAPI+ cells in the somatosensory cortex. (C) Absolute numbers of marker+ cells in the somatosensory cortex (SSC) and adjacent motor cortex (MC). (D) Representative confocal micrographs of immunohistochemistry with anti-Calbindin (CB, red) and anti-Somatostatin (SST, green) antibodies on WT, HET and KO tissue. Inserts: high magnification of SST+ cells. Nuclei were counterstained with DAPI (blue). (E) Quantification of the percentage of marker+ cells in each of 10 equal bins spanning the cortical wall. (F) Representative confocal micrographs of immunohistochemistry with anti-Parvalbumin (Pvalb, red) and anti-Calretinin (CR, green) antibodies on WT, HET and KO tissue. (G) Distribution of marker+ cells in each of 10 equal bins spanning the cortical wall, shown as percentage. All results are indicated as mean ± SEM. A minimum of 3 images per brain, obtained from four animals originating from three different litters were analyzed for each condition. *P < 0.05. Scale bars: 200 μm; insets: 50 μm.

### Altered behavior in *Pcdh19* mutant animals and their littermates

While correct cortical lamination is obviously important for proper brain function, the lack of any major lamination defects in the cortex of *Pcdh19* mutant animals does not preclude connectivity defects due to PCDH19 mosaicism that could become apparent at the behavioral level. As part of our characterization of the Taconic *Pcdh19* mutant mice, we also carried out a series of tests to determine whether these animals present any behavioral alterations. The paradigms included open field to evaluate general locomotor activity, anxiety and exploratory behavior, elevated plus maze to measure anxiety, and a social interaction test. We assessed animals at preweaning age and as adults, to account for any developmental effects. In addition to the WT littermates that *Pcdh19* mutant animals were housed with, we included a further control of single genotype housed WT animals (WT^SGH^) (Fig. 6A). Indeed, we note that a previous study on the X-linked ASD-related gene *Nlgn3*, also a membrane protein expressed in the developing cerebral cortex, revealed that housing conditions impact the behavior of wild-type animals when housed together with mutant animals (Kalbassi et al. 2017). The parents of the animals used to analyze behavior in the single genotype housed WT condition originated from our *Pcdh19* colony and behavior was analyzed separately for male and female mice.

**Figure 6.**
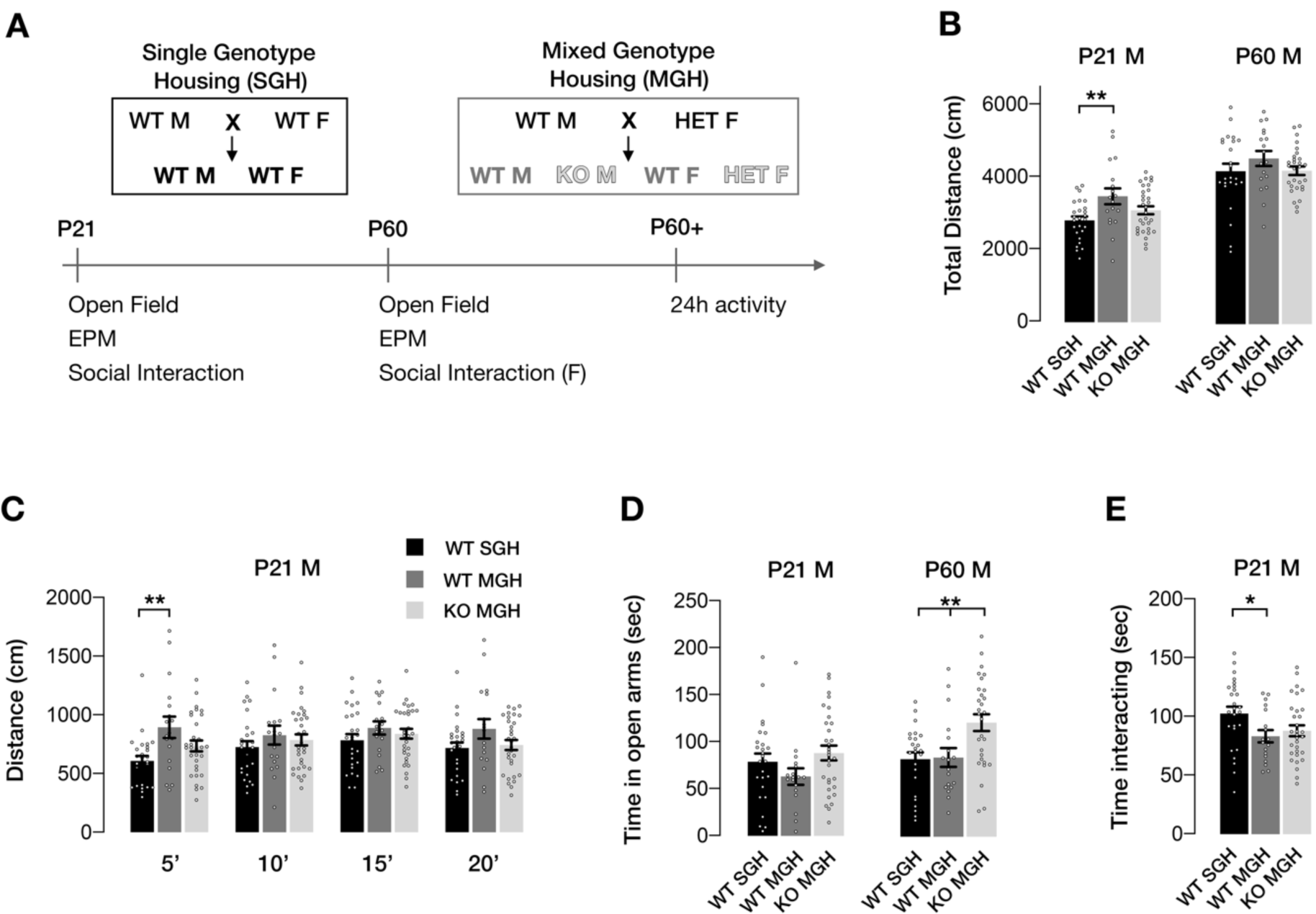
Behavioral alterations in *Pcdh19* KO males and their WT littermates. (A) Schematic of the behavioral experiments carried out. (B) Total distance travelled by males during the 20 minutes of the open field test at P21 and P60. (C) Distance travelled in the open field by P21 males, split in 5-minute intervals. (D) Total time spent by males in the open arms of the elevated plus maze during the 5-minute test at P21 and P60. (E) Time spent by P21 males interacting with a non-familiar female in oestrus. The total duration of the test was 5 minutes. Numbers of tested animals were: 26 WT^SGH^, 18 WT^MGH^, 31 KO at P21 and 24 WT^SGH^, 18 WT^MGH^, 29 KO at P60. Open field results correspond to the second day of testing at each age. Results are indicated as mean ± SEM, *P < 0.05; **P < 0.01. WT^SGH^, single genotype housed WT animals; WT^MGH^, mixed genotype housed animals.

Differences in male behavior were evident at P21 (Fig. 6B-E). Overall distance travelled during the 20 min open field paradigm was significantly different between genotypes (one way ANOVA, main effect of genotype *F*(2,72) = 5.02, *P* = 0.0091), with mixed genotype housed WT animals (WT^MGH^) showing a 23% increase compared to WT^SGH^ (Tukey: *q*(1,72) = 4.48, *P* = 0.0063; Fig. 6B). An analysis by 5-minute slots showed that the increased distance travelled by WT^MGH^ males was due to increased activity during the first 5 minutes (Kruskal-Wallis, *H* = 9.35, *P* = 0.0093, df = 2; post-hoc Dunn: *Z* = 3.01, *P* = 0.0079 WT^MGH^ vs WT^SGH^; Fig. 6C). This effect disappeared thereafter (Fig. 6C) and also when animals were tested again at ≥ P60 (Total distance: one way ANOVA, main effect of genotype *F*(2,68) = 1.13, *P* = 0.329; First 5 minutes: one way ANOVA, main effect of genotype *F*(2,68) = 1.31, *P* = 0.2759, Fig. 6B and Suppl. Fig. 5A). In accordance with these results, spontaneous activity (number of beam breaks) over a 24h period in adult male mice did not differ significantly between conditions (Suppl. Fig. 5B,C), neither when analyzed in total (one way ANOVA, main effect of genotype *F*(2,34) = 0.48, *P* = 0.621), nor in the light (one way ANOVA, main effect of genotype *F*(2,34) = 3.03, *P* = 0.0615) or dark periods (one way ANOVA, main effect of genotype *F*(2,34) = 0.31, *P* = 0.733). There were, however, differences at individual timepoints (19:00, 20:00, 8:00 and 10:00, Suppl. Fig. 5C). KO animals were less active during the start of the dark phase (19:00, Kruskal-Wallis, *H* = 16.08, *P* = 0.0003, df = 2; post-hoc Dunn: *Z* = 4.01, *P* = 0.0002 KO vs WT^MGH^; 20:00, one way ANOVA, main effect of genotype *F*(2,34) = 5.18, *P* = 0.0109; post hoc Tukey: *q*(1,34) = 4.42, *P* = 0.0099 HET vs KO vs WT^SGH^) and more active at 10:00 (10:00, Kruskal-Wallis, *H* = 10.78, *P* = 0.0046, df = 2; post-hoc Dunn: *Z* = 3.11, *P* = 0.0056 KO vs WT^MGH^; *Z* = 2.62, *P* = 0.0267 KO vs WT^SGH^). WT^MGH^ males were less active at 8:00 during the light period (8:00, Kruskal-Wallis, *H* = 7.17, *P* = 0.0277, df = 2; post-hoc Dunn: *Z* = 2.51, *P* = 0.0361 WT^MGH^ vs KO). Nevertheless, those differences do not seem to point to an overall activity defect and might be due to a smaller number of animals being tested.

To investigate whether the increased distance travelled by pre-weaned mixed genotype housed WT animals in the first 5 minutes of the open field could be due to increased anxiety, we analyzed the time spent in the center of the arena. No differences were found between the three conditions, neither at P21 (Kruskal-Wallis, *H* = 2.76, *P* = 0.2518, df = 2), nor at P60 (Kruskal-Wallis, *H* = 3.58, *P* = 0.1671, df = 2, Suppl. Fig. 5D). The results of the elevated plus maze confirmed the lack of differences at P21 (Kruskal-Wallis, *H* = 4.57, *P* = 0.1016, df = 2, Fig. 6D). However, this was not the case for adult animals, as adult KO males spent over 40% more time in the open arms than their WT^MGH^ littermates and WT^SGH^ controls, indicating reduced anxiety (one way ANOVA, main effect of genotype *F*(2,68) = 6.88, *P* = 0.0019; Tukey: *q*(1,68) = 4.10, *P* = 0.0138 KO vs WT^MGH^ and *q*(1,68) = 4.68, *P* = 0.0042 KO vs WT^SGH^; Fig. 6D).

Interestingly, we also detected a difference in social behavior at P21 (one way ANOVA, main effect of genotype *F*(2,72) = 3.39, *P* = 0.039, Fig. 6E). In this case, WT^MGH^ males spent significantly less time (19% decrease) interacting with an unfamiliar female in estrous than single-genotype housed WT males (Dunnett: *q*(1,72) = 2.37, *P* = 0.0382 WT^MGH^ vs WT^SGH^). KO males also showed a trend towards reduced interaction (14% decrease), but this difference was not significant (Dunnett: *q*(1,72) = 2.07, *P* = 0.0771 HET vs WT^SGH^).

Changes in behavior were more pronounced in female mice than in their male counterparts. We found again differences in the total distance travelled during the open field test at P21 (one way ANOVA, main effect of genotype *F*(2,69) = 9.54, *P* = 0.0002, Fig. 7A), with HET and WT^MGH^ females displaying an increase of 35% and 19%, respectively, when compared with single-genotype housed controls (Tukey: *q*(1,69) = 6.17, *P* = 0.0001 for HET vs WT^SGH^ and *q*(1,69) = 3.55, *P* = 0.0382 for WT^MGH^ vs WT^SGH^). Unlike in males, this effect was maintained at P60, but only in HET females, which travelled on average 19% more distance than WT^SGH^ animals (one way ANOVA, main effect of genotype *F*(2,69) = 3.99, *P* = 0.0229; Tukey: *q*(1,69) = 3.87, *P* = 0.0214 for HET vs WT^SGH^, Fig. 7A).

**Figure 7.**
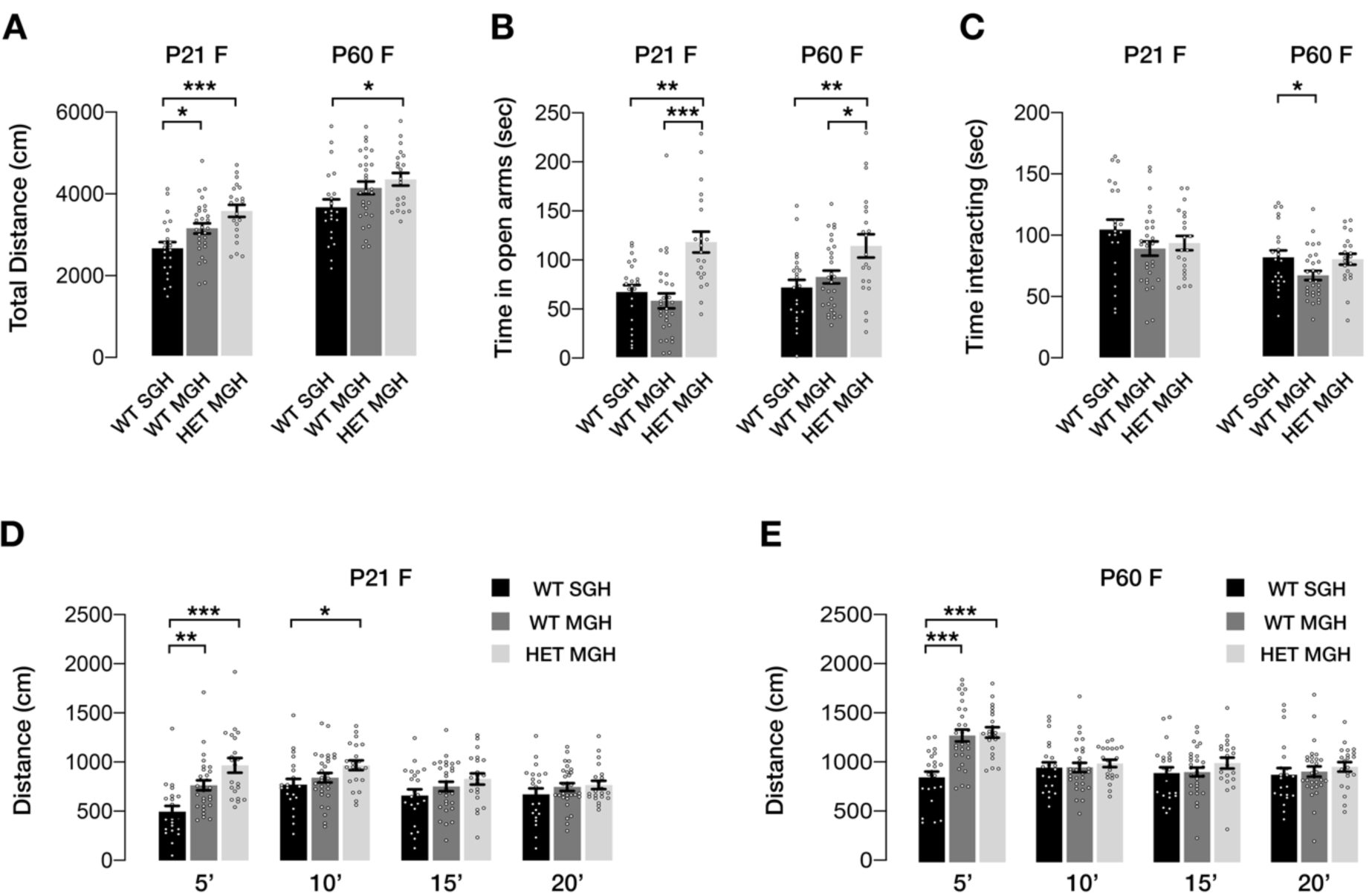
Behavioral alterations in *Pcdh19* HET females and their WT littermates. (A) Total distance travelled by females during the 20 minutes of the open field test at P21 and P60. (B) Total time spent by females in the open arms of the elevated plus maze during the 5-minute test at P21 and P60. (C) Time spent by females interacting with a non-familiar female at P21 and P60. The total duration of the test was 5 minutes. (D) Distance travelled in the open field by P21 females, split in 5-minute intervals. (E) Distance travelled in the open field by P60 females, split in 5-minute intervals. Numbers of tested animals were: 22 WT^SGH^, 29 WT^MGH^, 21 HET at P21 and P60. Open field results correspond to the second day of testing at each age. Results are indicated as mean ± SEM, *P < 0.05; **P < 0.01; ***P < 0.001. WT^SGH^, single genotype housed WT animals; WT^MGH^, mixed genotype housed animals.

Analysis by 5-minute intervals showed that the increase in total distance was mainly due to increased activity during the first 5 minutes in the open field arena both at preweaning age and in adults (P21: Kruskal-Wallis, *H* = 21.86, *P* < 0.0001, df = 2; P60: one way ANOVA, main effect of genotype *F*(2,69) = 17.95, *P* < 0.0001, Fig. 7D-E). This effect was significant at both ages for HET females and their WT siblings when compared with single-genotype housed females, with increases of 95% (HET) and 54% (WT^MGH^) at P21, and 53% (HET) and 49% (WT^MGH^) in adult animals (P21: Dunn: *Z* = 4.61, *P* < 0.0001 HET vs WT^SGH^ and *Z* = 3.12, *P* = 0.0055 WT^MGH^ vs WT^SGH^; P60: Tukey: *q*(1,69) = 7.38, *P* < 0.0001 HET vs WT^SGH^ and *q*(1,69) = 7.43, *P* < 0.0001 for WT^MGH^ vs WT^SGH^, Fig. 7D-E). HET females also travelled a significantly longer distance than WT^SGH^ females during the second 5-minute interval at P21, although the increase (25%) was much smaller in this case (one way ANOVA, main effect of genotype *F*(2,69) = 3.29, *P* = 0.0432; Tukey: *q*(1,69) = 3.58, *P* = 0.0359 HET vs WT^SGH^). Similarly to male mice, the spontaneous activity over 24h, measured as the number of beam breaks, was not altered for any of the three experimental groups in the light (one way ANOVA, main effect of genotype *F*(2,36) = 2.29, *P* = 0.1159), dark (one way ANOVA, main effect of genotype *F*(2,36) = 1.10, *P* = 0.3429) or total periods (one way ANOVA, main effect of genotype *F*(2,36) = 1.08, *P* = 0.3512) (Suppl. Fig. 6A,B). Again, isolated differences were evident at two timepoints during the dark phase (22:00, one way ANOVA, main effect of genotype *F*(2,36) = 3.84, *P* = 0.0309, Tukey: *q*(1,69) = 3.65, *P* = 0.0364 WT^MGH^ vs WT^SGH^; 4:00, Welch’s ANOVA *W*(2, 23.61) = 8.52, *P* = 0.0016, Dunnett T3: *t*(2, 18.32) = 3.83, *P* = 0.0036 WT^MGH^ vs HET; Dunnett T3: *t*(2, 23.41) = 2.65, *P* = 0.0417 WT^MHG^ vs WT^SGH^, Suppl. Fig. 6B), but no overall changes in activity were apparent in this test.

Since the increase in distance travelled during the first 5 minutes in the open field test does not seem to be caused by overall hyperactivity of HET animals and their WT siblings, we again analyzed anxiety-related behaviors in these animals. There were no differences in the time spent in the center of the open field arena for any of the conditions at P21 (Kruskal-Wallis, *H* = 4.68, *P* = 0.0962, df = 2) or P60 (Kruskal-Wallis, *H* = 4.09, *P* = 0.1296, df = 2; Suppl. Fig. 6C), but, similar to the results obtained with male animals, HET females spent significantly more time in the open arms of the elevated plus maze than any of the WT females at P21 (Kruskal-Wallis, *H* = 20.94, *P* < 0.0001, df = 2; Dunn: *Z* = 3.19, *P* = 0.042 between HET and WT^SGH^ and *Z* = 4.49, *P* < 0.0001 between WT^MGH^ and WT^SGH^) and P60 (one way ANOVA, main effect of genotype *F*(2,69) = 5.95, *P* = 0.0041; Tukey: *q*(1,69) = 4.67, *P* = 0.0043 for HET vs WT^SGH^ and *q*(1,69) = 3.72, *P* = 0.0281 between HET and WT^MGH^). The increases against WT^SGH^ and WT^MGH^ amounted to 76% and 103% at preweaning age and 60% and 39% in adults, indicating reduced anxiety, as seen also for adult male KO animals (Fig. 7B).

As in the case of male mice, the social interaction test revealed differences between single and mixed genotype housed WT females (Fig. 7C). However, this effect was only present in adult animals, with WT^MGH^ females spending 15% less time interacting with an unfamiliar female in estrous (One way ANOVA, main effect of genotype *F*(2,69) = 3.38, *P* = 0.0398; Dunnett: *q*(1,69) = 2.32, *P* = 0.0432 WT^MHG^ vs WT^SGH^).

Overall, we found significant behavioral differences between wild type and mutant animals that were generally more pronounced in HET females than in KO males. Importantly, we also uncovered an effect of housing on the behavior of WT animals.

## DISCUSSION

Although recent studies have shed light into the different functions of PCDH19 (Pederick et al. 2016; Hayashi et al. 2017; Pham et al. 2017; Bassani et al. 2018; Homan et al. 2018; Pederick et al. 2018, reviewed in Gerosa et al. 2019; Gécz and Thomas 2020), just exactly how mutations in this cell adhesion protein lead to the distinct symptoms associated with EIEE9 is still not well understood. Here we present a detailed analysis of neuronal sub-types expressing *Pcdh19/PCDH19* in the cortex of mice and humans. Our study reveals that *Pcdh19/PCDH19* is not only expressed in pyramidal neurons, but also in different types of interneurons, and that, in general, higher expression is limited to specific subpopulations in both cases. Our analysis also rules out major anomalies in the main axonal tracts and provides a quantitative assessment of cortical composition and lamination. Despite the lack of major lamination defects, out data suggest the possibility of subtle defects in layer composition that could contribute to the pathophysiology of EIEE9. Indeed, mutant animals display behavioral alterations in the open field (females) and elevated plus maze tests (males and females). Importantly, and as previously revealed with the analysis of *Nlgn3* mutants (Kalbassi et al. 2017), the *Pcdh19* mutation affects the behavior of wild-type littermates when housed in the same cage.

Hitherto, the characterization of the neuronal populations expressing PCDH19 has been hindered by the lack of specific antibodies that perform satisfactorily in immunohistochemistry analyses. In addition, as PCDH19 is likely distributed in both axons and dendrites (Pederick et al. 2016; Hayashi et al. 2017; Bassani et al. 2018), the unambiguous identification of cell bodies expressing PCDH19 is a challenging objective, as is the case for most membrane proteins in the cortex. To overcome this challenge, we focused on the expression of *Pcdh19* mRNA, which is detected in the cell soma and allows a better assessment of co-expression with other neuronal markers, which tend to be either nuclear or cytoplasmic. Although mRNA and protein expression are not necessarily correlated, available data show a good match between the regions with strongest mRNA and protein signals (Hayashi et al. 2017; Pederick et al. 2018). Our ISH/IHC combination approach provides experimental evidence for the expression of *Pcdh19* by different neuronal types across cortical layers, but it is inherently restricted to a particular brain region (in this case the SSC) and a few neuronal markers. However, the availability of datasets from scRNAseq studies, mainly carried out by the Allen Brain Atlas (https://portal.brain-map.org/atlases-and-data/rnaseq), allowed us to conduct a much more in depth expression analysis, not only in the mouse, but also in the human brain.

Our analysis of the mouse dataset published by Tasic et al in 2018 (Tasic et al. 2018) confirmed that *Pcdh19* is expressed by a majority of excitatory neurons in Layer V, projecting both intra- and extra-cortically, as well as by certain subtypes of Layer II/III projection neurons, in agreement with the ISH data. Expression in layer IV is minimal, and in layers VI and VIb it is mostly restricted to a couple of neuronal subtypes in the ALM. In interneurons, expression is widespread in the Sncg cluster, subtype specific in the Vip, Sst and Pvalb clusters, and very low in the Lamp5 and Serpinf1 clusters. These results demonstrate that while *Pcdh19* is expressed by a variety of excitatory and inhibitory neurons, expression remains specific for particular subtypes. This subtype specificity would suggest a role for PCDH19 in the establishment of neuronal circuits.

Human *PCDH19* follows a similar pattern, with expression in both excitatory and inhibitory neuronal types. We initially chose the MTG dataset to be able to establish a direct comparison between mouse and human neurons (Hodge et al. 2019). This comparison revealed shared high expression in Layer V excitatory neurons projecting through the pyramidal tract and in Chandelier cells. It also suggested that *Pcdh19* expression in mouse seems to be more widespread than in human, especially regarding excitatory neurons. However, *PCDH19* expression became much more apparent in the analysis of the Multiple Cortical Areas dataset that expanded the excitatory neuronal subtypes from 23 to 56. This probably reflects the splitting of subtypes that contained different *PCDH19* expressing populations into new subtypes in which *PCDH19* expressing cells cluster together. Nevertheless, expression in human excitatory neurons still seems to be more graded, with many more subtypes showing intermediate expression levels than in mouse. A similar picture emerges regarding expression in interneurons. High *PCDH19* expression can be found in subtypes of LAMP5, VIP, SST and PVALB interneurons, although most subtypes in humans seem to display some level of expression. One main difference between mouse and human, though, is the much lower expression of *PCDH19* in long range projecting interneurons in humans (Inh L3-6 SST NPY in the MTG dataset, Inh L6 SST NPY in the Multiple Cortical Areas dataset, Sst Chodl in the mouse dataset). The functional relevance of *Pcdh19*/*PCDH19* expression in particular neuronal subtypes will need to be established experimentally, but we expect our results to provide a framework to support those functional studies in the future, not least because of regional differences in the expression of this gene within neuronal subtypes.

Although two different *Pcdh19* KO models have been generated and published to date (Pederick et al. 2016; Hayashi et al. 2017), no detailed quantitative characterization of cortical composition and lamination has been reported. We have quantified 5 excitatory and 4 inhibitory markers, looking at overall abundance, as well as distribution throughout the cortical plate in the somatosensory cortex. In the case of parvalbumin, somatostatin and calretinin, the adjacent motor cortex was included in the study, due to the very low numbers of positive cells, whereas calbindin expressing neurons were much more abundant, especially in the upper layers, reflecting a population of excitatory neurons that also express this marker (van Brederode et al. 1991). Our analysis reveals no differences in the abundance of the different neuronal types and confirms the lack of major lamination defects (Pederick et al. 2016; Hayashi et al. 2017). However, our quantitative approach exposes more subtle changes in the distribution of certain neuronal types, suggesting the possibility of a slightly altered composition of particular layers or sublayers. The origin of these differences is unknown, but one possibility is that they could arise as a consequence of altered neurogenesis, since PCDH19 has been shown to play a role in this process (Fujitani et al. 2017; Homan et al. 2018; Lv et al. 2019). It is also important to consider that we carried out our analysis mainly in the SSC, but given that *Pcdh19* expression varies between cortical regions, it is possible that different areas might be affected in different ways by a total or partial loss of PCDH19. Reports of focal cortical dysplasia in EIEE9 patients (Kurian et al. 2018; Pederick et al. 2018) and focal areas of disorganization in ASD patients (Stoner et al. 2014) seem to support this possibility.

Despite the involvement of other delta protocadherins in the development of axonal tracts (Uemura et al. 2007; Piper et al. 2008; Biswas et al. 2014; Hayashi et al. 2014), our data do not support a major role of PCDH19 in this process. We did not detect any alterations in the main axonal tracts in the brain after staining for the axonal protein L1CAM, and a more detailed analysis of the corpus callosum also showed no differences in its dorso-ventral extension or the dorsal restriction of Neuropilin-1 expressing axons. This is in agreement with the lack of defects found by Hayashi et al in the projection of axons through this particular tract (Hayashi et al. 2017). More subtle defects in specific tracts would require much deeper analyses to be revealed.

Regardless of any anatomical alterations, assessing behavior allows a relevant functional assessment of the consequences of *Pcdh19* loss. Our analysis differed from the previously carried out with *Pcdh19* mutants (Hayashi et al. 2017; Lim et al. 2019). First, in addition to testing adult animals, we also tested animals at a much younger age (pre-weaning, P21), as EIEE9 is a developmental disorder and therefore it is relevant to determine when any behavioral changes begin. Second, we added a second cohort of control animals: wild-type single genotype housed mice, that have only been exposed to other WT animals during their life. An effect of WT littermates on the behavior of mutant animals was shown by Yang et al when they demonstrated that raising less sociable BTBR T+tf/J mice with highly sociable C57Bl6/J animals improved BTBR T+tf/J sociability (Yang et al. 2011). However, the impact of social environment on the behavior of WT littermates has only recently been demonstrated in a study with mice mutant for *Nlgn3*, an X-linked cell adhesion protein that has been implicated in ASD (Kalbassi et al. 2017). Therefore, this is further evidence to suggest that mutant mice can alter the behavior of their WT littermates.

In line with a previous study (Hayashi et al. 2017), changes in behavior were more apparent in HET females than in their KO male siblings. *Pcdh19* KO males only showed increased time spent in the open arms of the EPM, indicating reduced anxiety, when tested as adults. This same behavior was displayed by young *Pcdh19* HET females (P21), which maintained it into adulthood. However, HET females also exhibited increased exploratory behavior, or maybe hypersensitivity to new environments, from a young age, as demonstrated by their consistently higher distance travelled during the first 5 minutes in the open field at P21 and P60. It is important to consider that animals were placed into the open field arena 4 times in total, as they were tested on two consecutive days at both ages. Although habituation to the environment would be expected in this situation, the increased distance travelled during the first 5 minutes was apparent in all 4 trials, indicating a robust behavioral response. These results also suggest that behavioral changes in *Pcdh19* heterozygous animals start early in life, validating them as a good model for a developmental condition.

Open field and EPM tests were also performed in the study by Hayashi et al (Hayashi et al. 2017). They found no differences in the EPM, but whether this is due to experimental design or differences in the mouse model used is difficult to ascertain. Regarding the open field test, Hayashi et al found no differences in total distance or time in the center when the test was conducted at 11-12 weeks of age. However, when they repeated the test 23 weeks later, *Pcdh19* HET females spent significantly more time in the center of the open field arena, suggesting reduced anxiety. Although our animals did not display such behavior, they were tested around P60, which would be in agreement with the data from their first open field test. In addition, the results of our EPM test also indicate reduced anxiety in our animals, which could therefore represent a behavioral characteristic of *Pcdh19* mutant animals. Because no specific analysis of the first 5 minutes was carried out in that study, it is difficult to assess whether their animals exhibited increased activity during that period. Nevertheless, the fact that WT females display the same behavioral phenotype as their HET siblings indicates an effect of the social environment that can only be detected through the inclusion of single genotype housed WT animals. Interestingly, this effect was also present in young males, with WT^MGH^ travelling a higher distance in the first 5 minutes of the open field test than KO or WT^SGH^ males. However, unlike in the female population, this behavior disappeared in adulthood. Because adult male and female animals are housed separately, it is tempting to speculate that this effect of the social environment is somehow mediated by the HET females, although other causes, like a maternal effect, cannot be ruled out based on our experiments.

One of the comorbidities of EIEE9 patients is ASD (Kolc et al. 2020), and changes in *PCDH19* have also been linked to ASD cases (Piton et al. 2010; Harssel et al. 2013). Indeed, a recent behavioral study with the Taconic *Pcdh19* ko mouse model has revealed social interaction deficits in the three chamber test in KO males and HET females, as well as increased repetitive behavior in males (females were not tested) (Lim et al. 2019). In our analysis, we also found differences in social behavior, but, interestingly, only in WT^MGH^ animals. Both males and females spent less time interacting with a strange female at P21 and P60, respectively, than WT^SGH^ animals, in what appears to be another example of the effect of the environment on mouse behavior. Since males were not tested at P60, because at that age it becomes a measure of courtship behavior rather than simple social interaction and as such is not comparable to the P21 behavior, we don’t know if this phenotype would be maintained into adulthood or if, similar to the results of the open field, it would revert to normal with age. The fact that HET and KO animals did not differ in their behavior from their WT littermates is in contradiction with the results from Lim et al, although different tests were carried out in both studies, making a direct comparison difficult. In summary, our behavioral characterization of the *Pcdh19* Taconic mouse model reveals a stronger effect of *Pcdh19* mutation in HET females than in KO males and a significant effect of the social environment on the behavior of WT littermates, as previously described for *Nlgn3* mutant animals (Kalbassi et al. 2017). This effect should be taken into consideration for the design of future behavioral experiments, as failure to do so could result in misinterpretation of data and missed behavioral phenotypes.

## ACKNOWLEDGEMENTS

We would like to thank all members of the Martinez-Garay lab, as well as Y. Barde for insightful comments on this manuscript.

