## Supplementary material for "Cellular and behavioral characterization of *Pcdh19* mutant mice: subtle molecular changes, increased exploratory behavior and an impact of social environment": Suppl

### Supplementary Figure 1

## A

##### Excitatory neurons

| Mouse V1 & ALM clusters | Homology cluster | Human MTG clusters |
|---|---|---|
| L2/3 IT ALM Macc1 Lrg1<br>L2/3 IT ALM Sla<br>L2/3 IT ALM Ptrf<br>L2/3 IT VISp Rrad<br>L2/3 IT VISp Adamts2<br>L2/3 IT VISp Agmat | -Exc L2/3 IT- | Exc L2-3 LINC00507 GLP2R<br>Exc L2 LAMP5 LTK<br>Exc L2-3 LINC00507 FREM3 |
| L5 IT ALM Tmem163 Dmrtb1<br>L5 IT VISp Hsd11b1 Endou | -Exc L3/5 IT- | Exc L3-4 RORB CARM1P1 |
| L5 IT ALM Npw<br>L5 IT ALM Cbln4 Fezf2<br>L5 IT ALM Pdl5<br>L5 IT ALM Lypd Gpr88<br>L4 IT VISp Rspo1 | -Exc L4/5 IT- | Exc L3-5 RORB FILIP1L<br>Exc L3-5 RORB TWIST2<br>Exc L3-5 RORB ESR1 |
| L5 PT ALM Hpgd<br>L5 PT ALM Npsr1<br>L5 PT VISp Lgr5<br>L5 PT VISp C1ql2 Cdh13<br>L5 PT VISp C1ql2 Ptgfr<br>L5 PT VISp Krt80 | -Exc L5 PT- | Exc L4-5 FEZF2 SCN4B |
| L5 IT ALM Tnc<br>L6 IT VISp Col18a1 | -Exc L5/6 IT 1- | Exc L3-5 RORB COL22A1 |
| L5 IT ALM Tmem163 Arhgap25<br>L5 IT ALM Cpa6 Gpr88<br>L5 IT VISp Whrn Tox2<br>L5 IT VISp Batf3 | -Exc L5/6 IT 2- | Exc L4-5 RORB DAPK2<br>Exc L4-6 RORB SEMA3E<br>Exc L4-5 RORB FOLH1B |
| L5 IT VISp Col6a1 Fezf2<br>L6 IT ALM Tgfb1<br>L5 IT VISp Col27a1 | -Exc L5/6 IT 3- | Exc L4-6 RORB C1R<br>Exc L5-6 RORB TTC12 |
| L6 NP ALM Trh<br>L5 NP ALM Trhr Nefl<br>L5 NP VISp Trhr Met<br>L5 NP VISp Trhr Cpne7 | -Exc L5/6 NP- | Exc L4-6 FEZF2 IL26 |
| L6 CT ALM Cpa6<br>L6 CT VISp Gpr139<br>L6 CT VISp Krt80 Sla<br>L6 CT VISp Ctxn3 Sla<br>L6 CT VISp Nxph2 Wls<br>L6 CT VISp Ctxn3 Brinp3 | -Exc L6 CT- | Exc L5-6 FEZF2 ABO |
| L6 IT VISp Penk Fst<br>L6 IT VISp Col23a1 Adamts2<br>L6 IT VISp Penk Col27a1 | -Exc L6 IT 1- | Exc L5-6 THEMIS C1QL3 |
| L6 IT ALM Oprk1<br>L6 IT VISp Car3 | -Exc L6 IT 2- | Exc L5-6 THEMIS DCSTAMP<br>Exc L5-6 THEMIS FGF10<br>Exc L5-6 THEMIS CRABP1 |
| L6b ALM Olfr111 Spon1<br>L6b ALM Olfr111 Nxph1<br>L6b VISp Crh<br>L6b VISp Col8a1 Rxfp1<br>L6b Hsd17b2<br>L6b VISp Mup5<br>L6b P2ry12 | -Exc L6b- | Exc L6 FEZF2 OR2T8<br>Exc L6 FEZF2 SCUBE1<br>Exc L5-6 FEZF2 EFTUD1P1 |

## B

##### Interneurons

| Mouse V1 & ALM clusters | Homology cluster | Human MTG clusters |
|---|---|---|
| Pvalb Vipr2 | -Chandelier- | Inh L2-5 PVALB SCUBE3 |
| Lamp5 Plch2 Dock5 | -Lamp5 1- | Inh L1-2 LAMP5 DBP |
| Lamp5 Fam19a1 Pax6<br>Lamp5 Fam19a1 Tmem182<br>Lamp5 Ntn1 Npy2r | -Lamp5 2- | Inh L1 SST CHRNA4 |
| Lamp5 Lhx6 | -Lamp5 Lhx6- | Inh L2-6 LAMP5 CA1 |
| Lamp5 Lsp1 | -Lamp5 Rosehip- | Inh L1-4 LAMP5 LCP2 |
| Lamp5 Krt73 | -Pax6- | Inh L1-2 PAX6 TNFAIP8L3<br>Inh L1-2 PAX6 CDH12 |
| Pvalb Gpr149 Islr<br>Palb Akr1c18 Ntf3<br>Sst Nts<br>Pvalb Th Sst<br>Pvalb Gabrg1 | -Pvalb 1- | Inh L5-6 PVALB LGR5<br>Inh L5-6 SST TH<br>Inh L4-5 PVALB MEPE<br>Inh L5-6 SST MIR548F2 |
| Pvalb Sema3e Kank4<br>Palb Calb1 Sst<br>Pvalb Reln Tac1<br>Pvalb Reln Itm2a<br>Pvalb Tpbg | -Pvalb 2- | Inh L2-4 PVALB WFDC2<br>Inh L4-6 PVALB SULF1 |
| Sst Myh8 Fibin<br>Sst Chrna2 Glra3<br>Sst Myh8 Etv1<br>Sst Nr2f2 Necab1<br>Sst Chrna2 Ptgdr | -Sst 1- | Inh L3-6 SST HPGD<br>Inh L4-6 SST B3GAT2 |
| Sst Crhr2 Efemp1<br>Sst Chr 4930553C11Rik<br>Sst Esm1 | -Sst 2- | Inh L5-6 SST KLHDC8A |
| Sst Tac2 Tacstd2<br>Sst Rxfp1 Eya1<br>Sst Rxfp1 Prdm8 | -Sst 3- | Inh L4-6 SST GXYLT2<br>Inh L5-6 SST NPM1P10 |
| Sst Hpse Cbln4<br>Sst Hpse Sema3c | -Sst 4- | Inh L3-5 SST ADGRG6<br>Inh L4-5 SST STK32A |
| Sst Calb2 Necab1<br>Sst Tac1 Tacr3<br>Sst Tac1 Htr1d<br>Sst Calb2 Pdlim5<br>Sst Mme Fam114a1 | -Sst 5- | Inh L1-3 SST CALB1 |
| Sst Chodl | -Sst Chodl- | Inh L3-6 SST NPY |
| Vip Igfbp6 Pltp<br>Vip Igfbp6 Car10 | -Vip 1- | Inh L1-4 VIP PENK<br>Inh L1-3 VIP ADAMTSL1<br>Inh L1-2 SST BAGE2 |
| Vip Lmo1 Fam159b<br>Vip Arhgap36 Hmcn1<br>Vip Lmo1 Myl1<br>Vip Gpc3 Slc18a3 | -Vip 2- | Inh L2-6 VIP QPCT<br>Inh L3-6 VIP HS3ST3A1 |
| Serpinf Aqp5 Vip<br>Vip Pygm C1ql1<br>Vip Chat Htr1f | -Vip 3- | Inh L2-5 VIP SERPINF1<br>Inh L2-5 VIP TYR<br>Inh L1-2 VIP PCDH20 |
| Vip Rspo1 Itga4<br>Vip Lect1 Oxtr<br>Vip Rspo4 Rxfp1 Chat<br>Vip Ptprt Pkp2 | -Vip 4- | Inh L2-4 VIP CBLN1<br>Inh L1-3 VIP CCDC184<br>Inh L1-3 VIP GGH<br>Inh L1-3 VIP CHRM2 |
| Vip Crispld2 Htr2c<br>Vip Crispld2 Kcne4 | -Vip 5- | Inh L2-4 VIP CHRNA6<br>Inh L2-4 VIP SPAG17<br>Inh L1-2 VIP LBH<br>Inh L1-4 VIP OPRM1<br>Inh L2-3 VIP CASC6 |
| Sncg Gpr50<br>Serpinf1 Clrn1<br>Sncg Vip Nptx2<br>Sncg Slc17a8<br>Sncg Vip Itih5<br>Scng Col15a1 Pde1a | -Vip Scng- | Inh L1-2 VIP TSPAN12 |

**Supplementary Figure 1**

Diagram showing the homology clusters defined by Hodge et al and the corresponding mouse and human neuronal subtypes assigned to each cluster. (A) Excitatory neuronal clusters. (B) Inhibitory neuronal clusters.

### Supplementary Figure 2

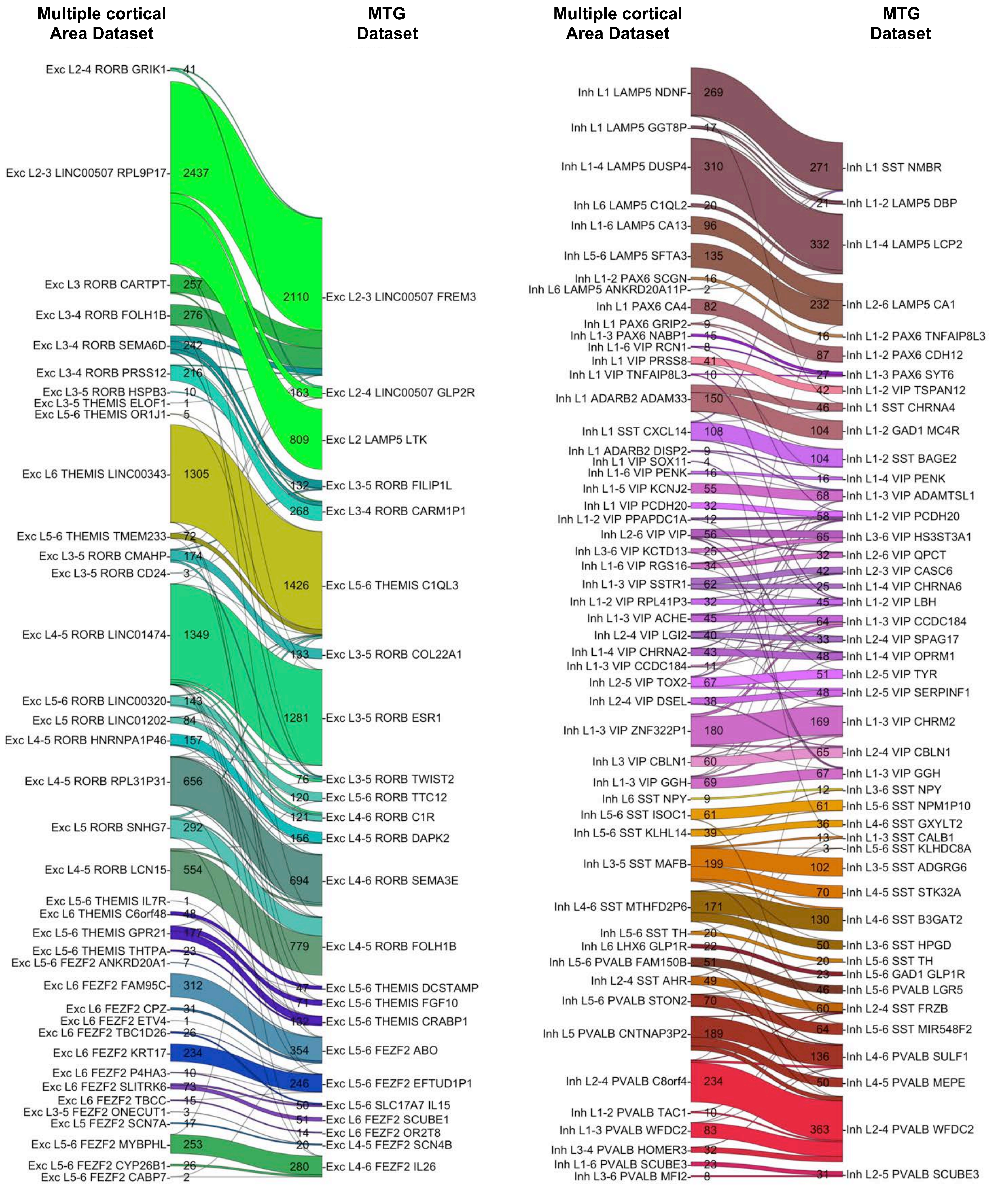


**Supplementary Figure 2**

River plots showing the mapping of the nuclei from the MTG dataset to the subtypes defined by the Multiple Cortical Areas dataset, containing nuclei from several different brain regions. (A) Mapping of excitatory neurons. (B) Mapping of inhibitory neurons.

#### Supplementary Figure 3

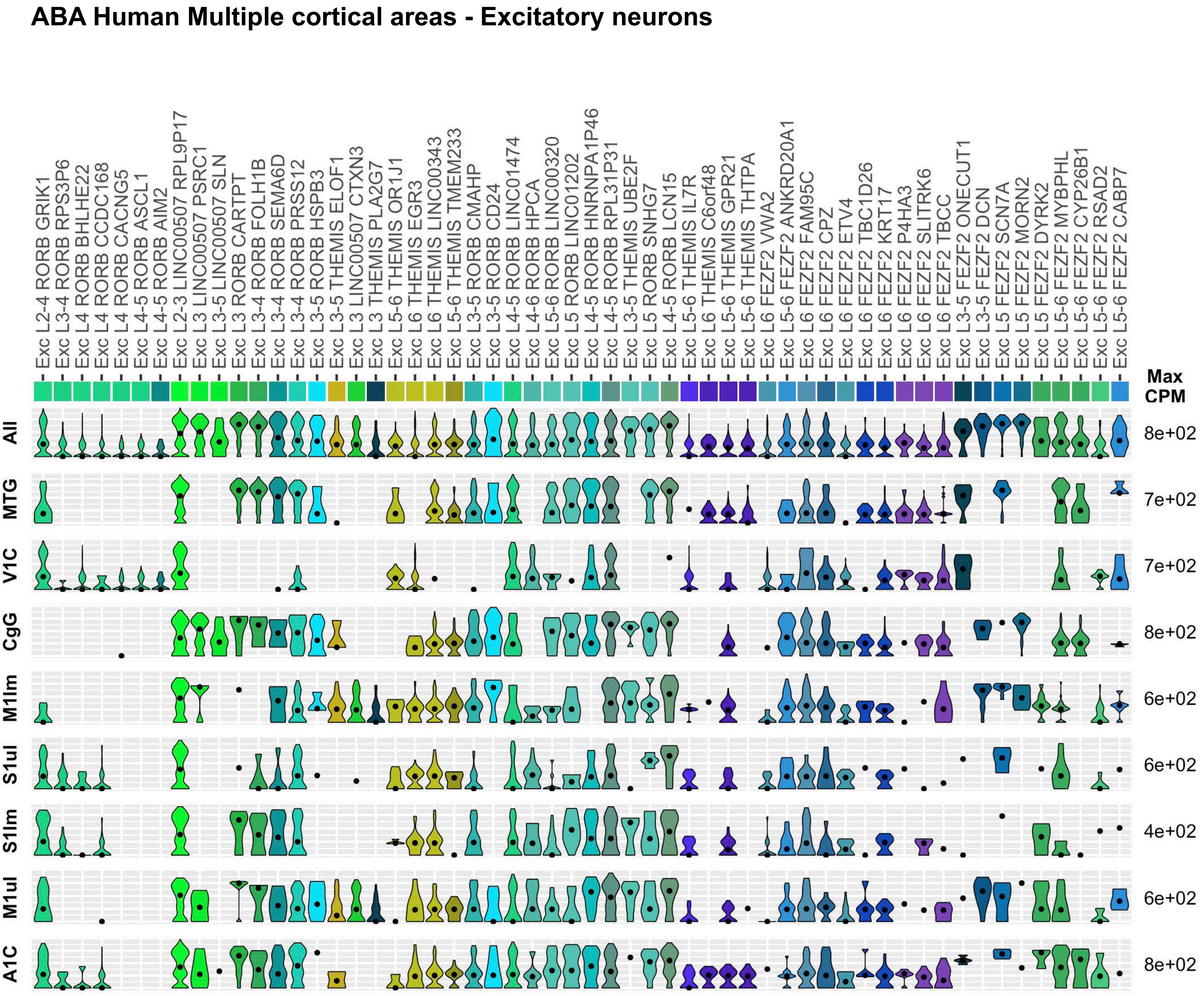


**Supplementary Figure 3**

Gene expression and distribution of *PCDH19* in cortical excitatory projection neurons of the Allen Brain Atlas human Multiple Cortical Areas dataset, represented by violin plots. The first row shows overall expression of *PCDH19* in the combined dataset, subsequent rows show expression by brain region. Dots indicate the median value of the population. Absence of a violin plot in a row indicates no cells from that particular brain region were mapped to the corresponding neuronal subtype. MTG, middle temporal gyrus; V1C, primary visual cortex; CgG, anterior cingulate gyrus; M1lm, primary motor cortex, lower limb region; S1ul primary somatosensory cortex upper limb region; S1lm, primary somatosensory cortex lower limb region; M1ul primary motor cortex, upper limb region, A1C, primary auditory cortex.

#### Supplementary Figure 4

##### ABA Human Multiple cortical areas - Interneurons

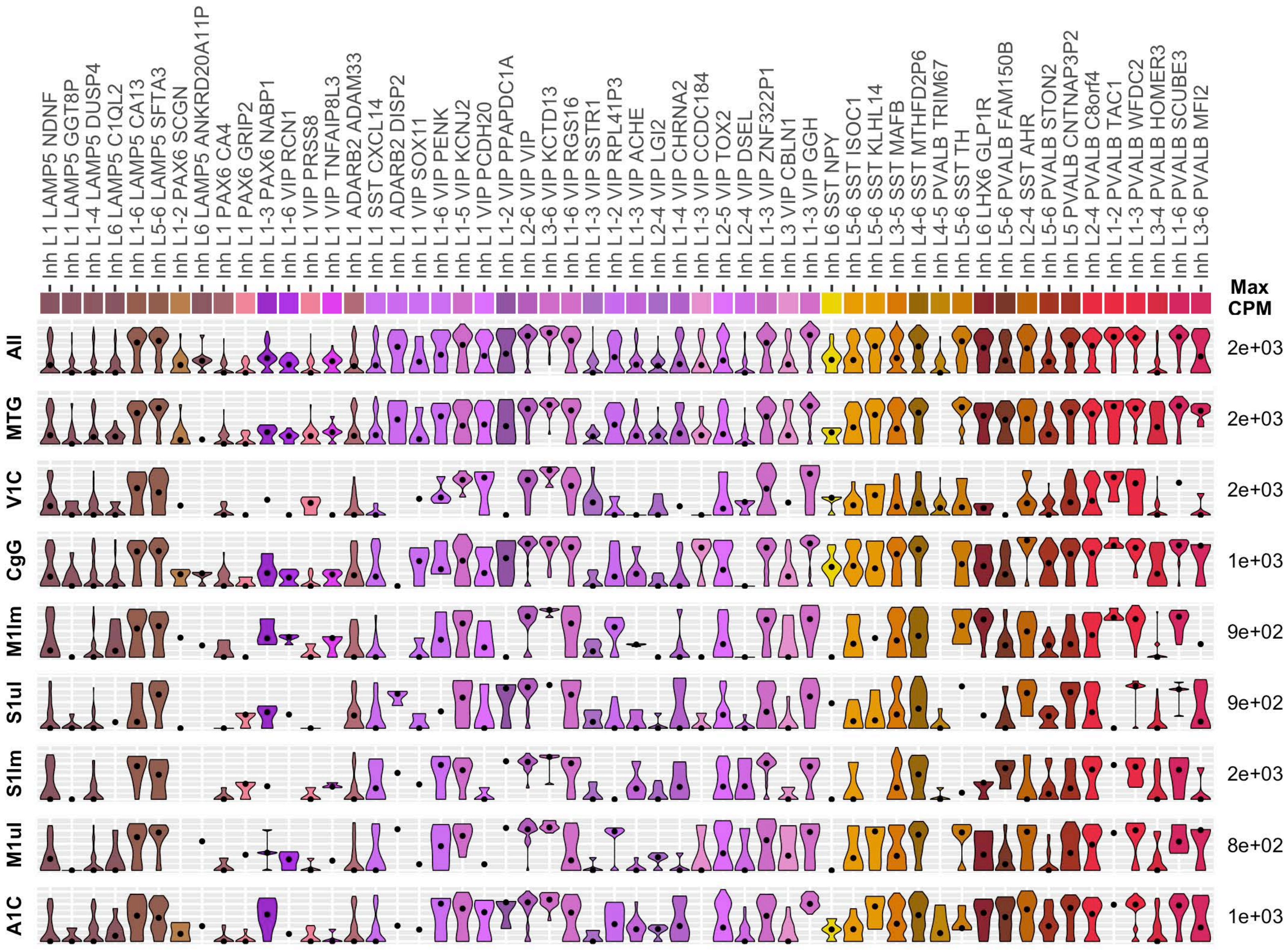


**Supplementary Figure 4**

Gene expression and distribution of *PCDH19* in cortical inhibitory neurons of the Allen Brain Atlas human Multiple Cortical Areas dataset, represented by violin plots. The first row shows overall expression of *PCDH19* in the combined dataset, subsequent rows show expression by brain region. Dots indicate the median value of the population. Absence of a violin plot in a row indicates no cells from that particular brain region were mapped to the corresponding neuronal subtype. MTG, middle temporal gyrus; V1C, primary visual cortex; CgG, anterior cingulate gyrus; M1lm, primary motor cortex, lower limb region; S1ul primary somatosensory cortex upper limb region; S1lm, primary somatosensory cortex lower limb region; M1ul primary motor cortex, upper limb region, A1C, primary auditory cortex.

#### Supplementary Figure 5

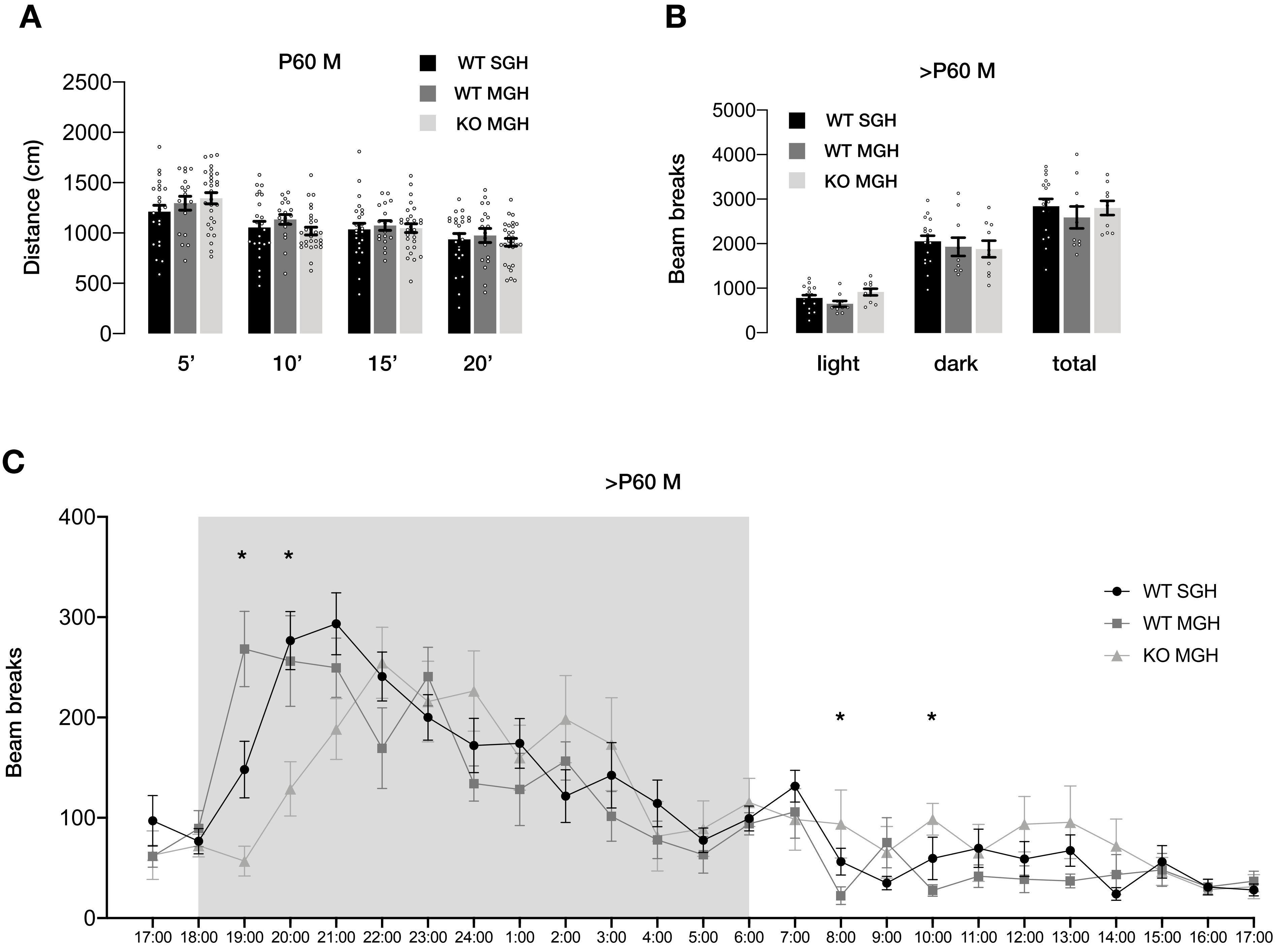


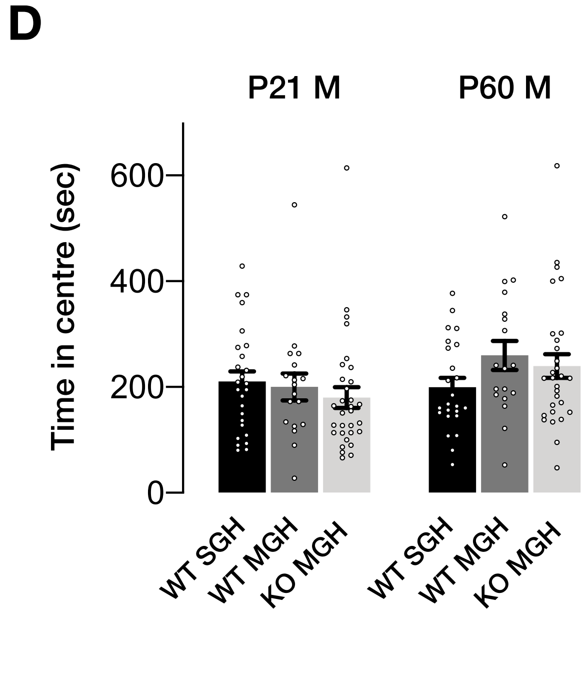


**Supplementary Figure 5**

(A) Distance travelled in the open field by P60 males, split in 5-minute intervals. (B) Number of beam breaks during the 24-hour activity test for the males of the different conditions. Light: light phase, dark: dark phase. (C) Number of beam breaks by hour in the 24-hour activity test for the males of the different conditions. The time of the day is shown on the X-axis and the grey square indicates the hours of the dark period. (D) Time spent by males in the center of the arena during the 20-minutes open field test at P21 and P60. Numbers of tested animals were: 26 $WT^{SGH}$, 18 $WT^{MGH}$, 31 KO at P21 and 24 $WT^{SGH}$, 18 $WT^{MGH}$, 29 KO at P60. For the 24-hour activity test, number were 17 $WT^{SGH}$, 10 $WT^{MGH}$ and 10 KO. Results are indicated as mean ± SEM. $^{*}P < 0.05$. $WT^{SGH}$, single genotype housed WT animals; $WT^{MGH}$, mixed genotype housed animals.

#### Supplementary Figure 6

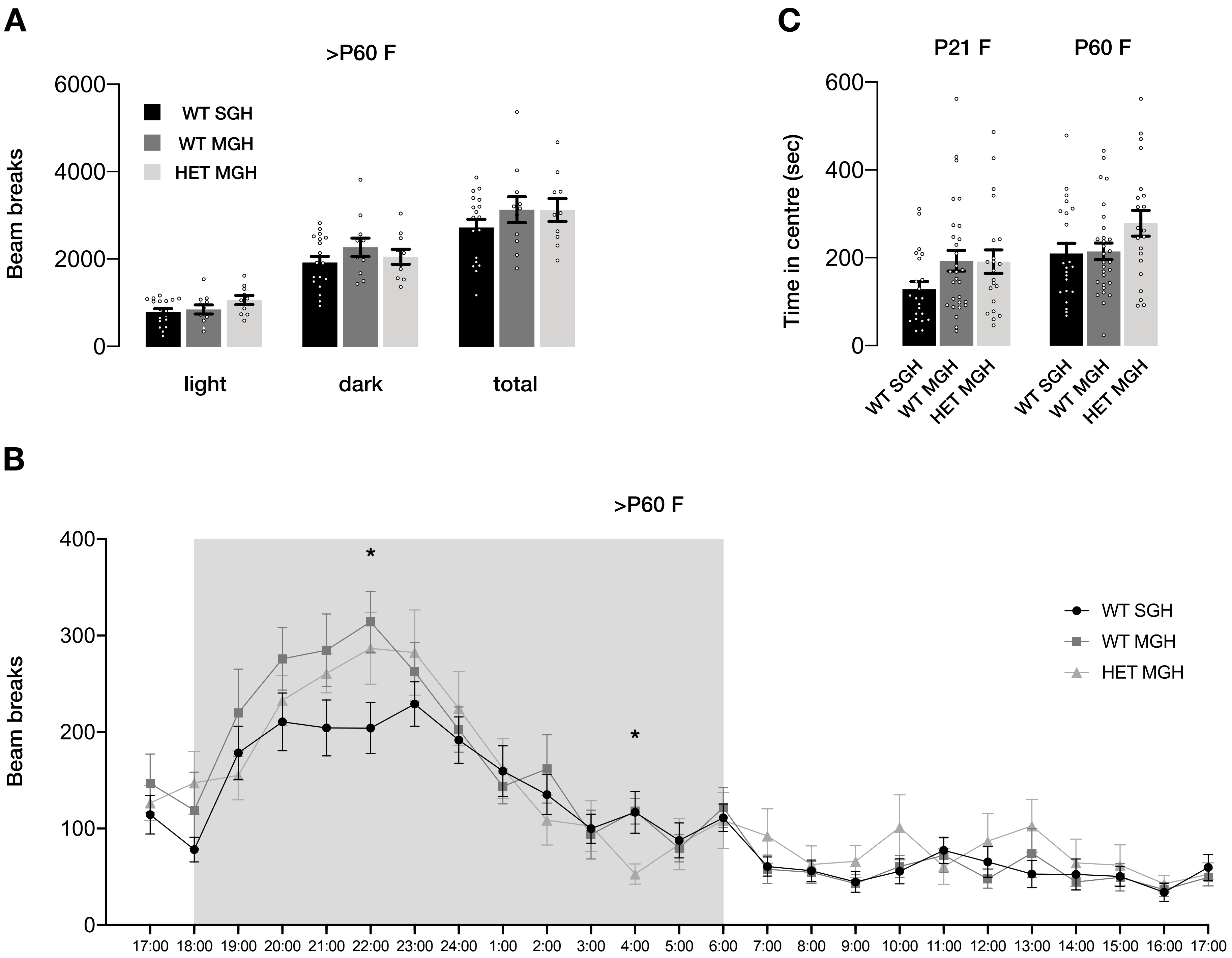


**Supplementary Figure 6**

(A) Number of beam breaks during the 24-hour activity test for the females of the different conditions. Light: light phase, dark: dark phase. (CB Number of beam breaks by hour in the 24-hour activity test for the females of the different conditions. The time of the day is shown on the X-axis and the grey square indicates the hours of the dark period. (C) Time spent by females in the center of the arena during the 20-minutes open field test at P21 and P60. Numbers of tested animals were: 22 $WT^{SGH}$, 29 $WT^{MGH}$, 21 HET at P21 and P60. For the 24-hour activity test, number were 18 $WT^{SGH}$, 11 $WT^{MGH}$ and 10 HET. Results are indicated as mean ± SEM. *P < 0.05. $WT^{SGH}$, single genotype housed WT animals; $WT^{MGH}$, mixed genotype housed animals.
